# Pollutant biodegradation profile mediated by multi-trophic microbial dynamics in rivers

**DOI:** 10.64898/2026.01.14.699447

**Authors:** Joeselle M. Serrana, Run Tian, Francisco J. A. Nascimento, Elias Broman, Benoît Dessirier, Malte Posselt

## Abstract

Microbial communities and environmental conditions are closely linked to ecosystem functions and directly govern the biodegradation of pollutants in aquatic environments. However, the role of multi-trophic interactions and their spatiotemporal dynamics in these processes remains poorly understood. Here, we examined how seasonal and spatial variations, mediated by trophic interactions within benthic microbial communities, influence their composition, functional capacity, and collective potential to degrade a diverse array of organic pollutants in rivers. By characterizing both prokaryotic (i.e., archaea and bacteria) and eukaryotic taxa (i.e., algae, fungi, protists, and metazoans), and inferring metabolic pathways, we explored the connections between community composition and pollutant degradation in wastewater-receiving rivers across four seasons. Mediation analysis revealed that multi-trophic communities mediate the total effect of environmental factors on the biodegradation of 96 organic pollutants. Prokaryotic communities explained 60% of the total environmental influence on pollutant biodegradation. Additionally, eukaryotic groups had significant indirect mediation effects, with fungal, protistan, algal, and metazoan communities responsible for 56%, 53%, 26%, and 38%, respectively. Notably, fungal and protist communities mediated approximately 83% and 73% of the environmental impacts on prokaryotic community composition, respectively. Across the two rivers studied, spatial variation (at the river and reach scales) explained more variance in community composition than seasonality over the sampled year. Our findings improve understanding of ecosystem resilience and support the development of predictive models and sustainable water management strategies in dynamic aquatic environments.

## Introduction

Rivers provide a wide range of ecosystem services, including water purification and waste treatment, driven by interconnected physical, chemical, and biological processes [1, 2]. These services are sustained by dynamic interactions within river networks, ranging from hydrological regimes to biotic communities, that regulate nutrient cycling and organic matter decomposition [3]. Among the biological drivers, microorganisms play a pivotal role in transforming nutrients, degrading pollutants, and neutralizing harmful substances through metabolic activity, effectively mitigating their negative ecological impacts [4]. Hence, microbial communities play a crucial role in the natural purification functions of river ecosystems, making significant contributions to environmental health and sustainable water management [5, 6].

Several studies have explored the relationship between microbial composition and the degradation of pollutants, e.g., pesticides, pharmaceuticals, and industrial chemicals, in aquatic systems [7–9]. Experimental flume studies have further demonstrated that bacterial diversity influences the transformation of wastewater-derived compounds [10–12], with recent findings showing that bacterial community dynamics are highly sensitive to complex pollutant mixtures [7, 13]. While much of this research has focused on prokaryotes, emerging evidence highlights the role of microbial eukaryotes in pollutant degradation. Eukaryotic taxa, including protists, fungi, and algae, contribute to the ecosystem through enzymatic breakdown, nutrient cycling, and trophic interactions [e.g., 14–17]. Protists, for instance, regulate bacterial populations via predation, indirectly shaping microbial community structure and function [18–20]. Fungi and algae also participate in organic matter decomposition and nutrient assimilation, often synergistically with prokaryotes [21]. Additionally, interactions between sediment-dwelling metazoans (e.g., nematodes and ostracods) and bacteria have been shown to influence pollutant transformation processes [22]. Despite these insights, the ecological mechanisms by which trophic interactions among prokaryotes and microbial eukaryotes influence pollutant biodegradation remain poorly understood. Addressing this gap requires a multi-trophic perspective, one that integrates species interactions across trophic levels to understand how ecological complexity governs ecosystem functions, i.e., biodegradation, in aquatic systems [23, 24].

Moreover, the extent to which environmental factors modulate these interactions and their collective biodegradation capacity is still unclear. River ecosystems are inherently dynamic, with spatial and temporal variations in temperature, nutrient availability, and flow regimes shaping community composition and function across trophic levels [17, 25, 26]. For example, previous studies have documented how river geomorphology, hydrology, and anthropogenic inputs influence microbial diversity across river segments [27, 28]. Seasonal changes further drive transient shifts in microbial processes across spatial scales [29, 30]. For example, Tian et al. [31] reported seasonal variability in pollutant degradation rates, with faster transformation in warmer months for many compounds. Interestingly, these patterns deviated from classical temperature-dependent models, suggesting adaptive shifts in microbial function and further highlighting the need to understand how seasonal and spatial factors interact with community dynamics. This growing body of evidence underscores the need to examine how multi-trophic communities mediate the influence of environmental variation on pollutant degradation. Understanding these dynamics could be key to predicting ecosystem resilience and the efficiency of microbial processes in rivers.

Using benthic sediment samples and chemical analytical data collected by Tian et al. [31], we hypothesize that (i) seasonal and spatial factors significantly shape the composition and functional capacity of riverine microbial communities, and (ii) that trophic interactions within these communities mediate the influence of environmental factors on pollutant degradation by regulating microbial composition and driving transformation pathways. To test these hypotheses, we profiled benthic prokaryotes and eukaryotes (inc. algae, fungi, protists, and metazoans) across the upstream and downstream reaches of two rivers over four seasons and inferred the prokaryotes’ metabolic pathways to assess spatiotemporal shifts in community structure and function. We then examined how these communities relate to the rivers’ capacity to degrade a complex mixture of 96 organic pollutants across diverse use categories (e.g., agrochemicals, cosmetics, industrial chemicals, and pharmaceuticals).

## Materials and methods

### Sample description

Tian et al. [31] collected benthic sediments and environmental samples in 2022 from two small rivers, i.e., Vitsån and Knivstaån, near Stockholm, Sweden (**Figure 1A**), at two reaches, i.e., one upstream and one downstream of the rivers receiving municipal wastewater. The environmental parameters recorded in the field included pH, dissolved oxygen, electrical conductivity, and water temperature (**Supplementary Table S1** and **Figure S1**). For detailed site and sampling information, please refer to Tian et al. [31]. Additional parameters and water chemistry data, e.g., flow rate, total organic carbon (TOC), nutrients, total suspended solids (TSS), and heavy metal concentrations, were aggregated based on modeled data for Vitsån and Knivstaån from a national runoff model (HYPE) and on time series collected at proxy sites in Vitsån and Knivstaån by the national environmental monitoring program [32] (**Supplementary Text S1** and **Table S1**). Environmental parameters, except pH, were log-transformed prior to statistical analysis. The samples were also grouped into different spatiotemporal categories, i.e., spatial (by River, Reach, or River × Reach) or temporal (by Season, **Supplementary Text S1**), as well as their combinations (e.g., River × Reach × Season).

**Figure 1.**
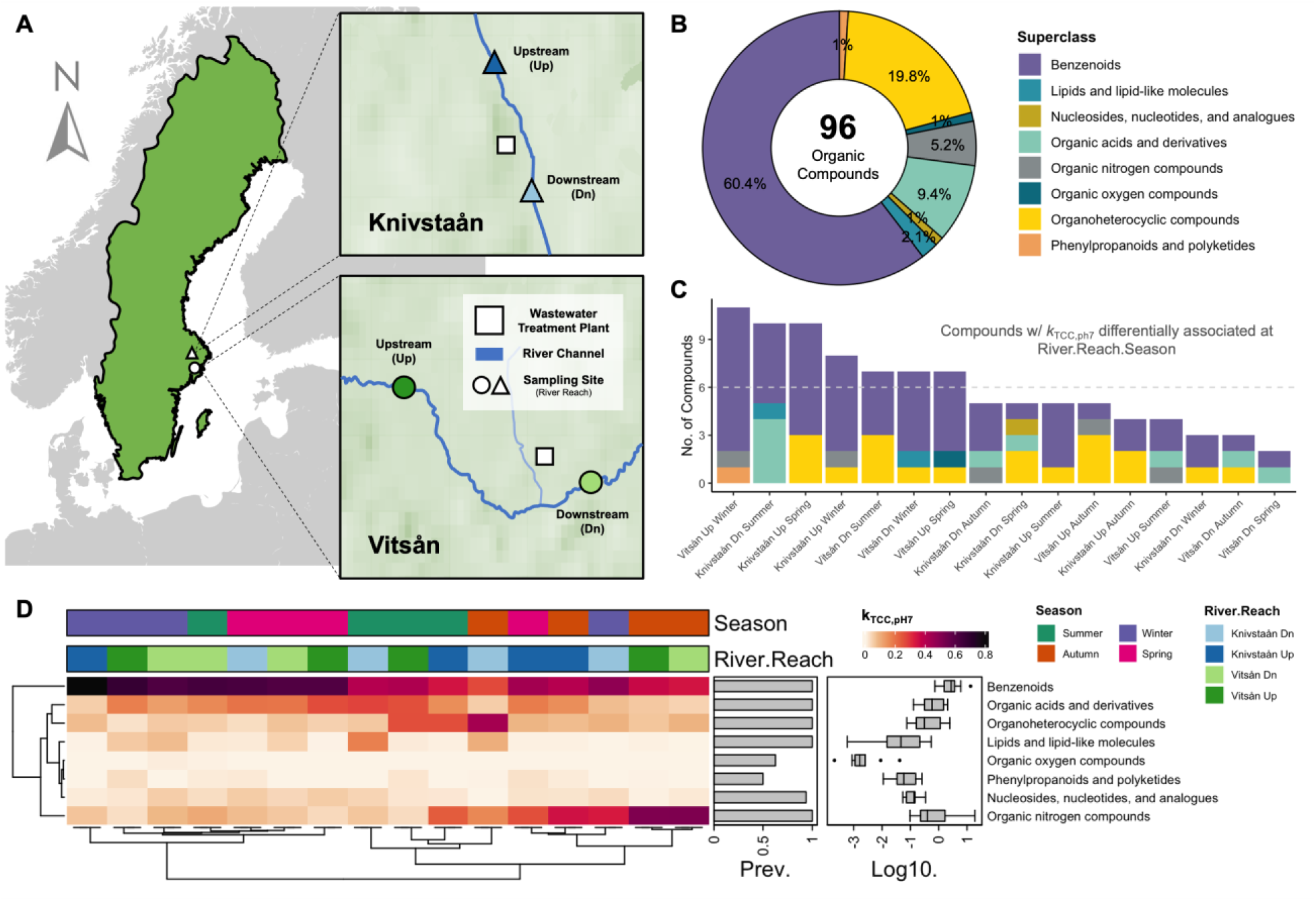
Influence of spatiotemporal variation on the biodegradation profile of organic pollutants in rivers (Tian et al., 2024). (A) Map showing the sampling sites of the two rivers in Sweden assessed in the study. (B) Relative representation of the 96 organic compounds used to profile the biodegradation potential of the rivers. The superclass category is based on structure-based chemical taxonomy. (C) Frequency of the organic compounds with biodegradation rates (k_TCC,pH7_) that are differentially associated at the spatiotemporal (River × Reach × Season) grouping. Differential abundance analysis based on a random forest test on the biodegradation profile of the organic compounds. (D) Spatiotemporal pattern of the biodegradation rates grouped at the superclass level across the rivers and their reaches. Hierarchical clustering analysis based on Euclidean distance.

### Biodegradation experiment and chemical analytics

The biodegradation experiments were conducted in accordance with a modified OECD 309 test guideline [33]. Four experiments were conducted in 10-day incubation setups, with each seasonal sample tested based on the river water temperatures at the time of sampling for environmental relevance (i.e., summer at 19°C, autumn at 11°C, winter at 4°C, and spring at 17°C). Sediment samples from before and after the experiment were collected and stored at −80°C for DNA extraction. Chemical analysis was conducted using a UHPLC-Orbitrap-MS/MS (Thermo Fisher Scientific, San Jose, CA) with electrospray ionization, and mass spectral data were processed with Compound Discoverer 3.3 software. Details of the biodegradation experiment and chemical analytics are available in the published study [31]. The pH 7 and total cell count (TCC) corrected first-order rate constants (k_TCC,pH7_) were used in the downstream analysis (**Supplementary Table S2**). Negative rate constants (15% of the data) were considered artifacts from the sorption correction and were excluded from the biodegradation profile. The compounds were assigned into structure-based chemical taxonomic categories (ChemOnt, e.g., superclasses, classes, and parent levels) using the ClassyFire Batch tool [34] (**Figure 1B-C** and **Supplementary Table S2**).

### Amplicon sequencing and bioinformatics

Total genomic DNA was extracted from the sediment samples and then amplified for prokaryotic 16S rRNA V4-V5 region using the 515F-907R primers, and eukaryotic 18S rRNA V4 region using the 528F-706R primers, followed by high-throughput sequencing on the Illumina NovaSeq 6000 platform, generating 250 bp paired-end reads. The raw sequence reads were then processed using the DADA2 v.1.32.0 [35] pipeline to filter noise, remove chimeras, and infer amplicon sequence variants (ASVs), which were taxonomically assigned using the RDP naive Bayesian classifier method [36] against the Silva v.138.2 [37] database for 16S rRNA and the Protist Ribosomal Reference (PR2) v.5.1.0 [38] database for 18S rRNA data. The datasets were then assigned to five categories: 16S data for prokaryotes (bacteria and archaea) and 18S rRNA data for algae, fungi, protists, or metazoans. Putative functional pathways were predicted from the 16S rRNA data to the KEGG database [39] using the Tax4Fun2 v.1.1.5 tool [40]. More details regarding amplicon sequencing and bioinformatics are presented in the **Supplementary Text S2** and **Tables S3-S7**.

### Statistical analyses

All visualization and statistical analysis were conducted using R v.4.4.2 [41]. After assessing differences in microbial profiles between the before- and after-incubation sediment samples, the before-incubation sample was used to represent each sample’s microbial community profile in subsequent analyses (**Supplementary Text S3**). Taxonomic and phylogenetic alpha diversity indices were estimated, and the beta diversity of the benthic microbiome was assessed using weighted UniFrac distance and Bray-Curtis dissimilarity indices. Principal coordinate analysis (PCoA) was performed to visualize the differences in community composition between groups. An analysis of similarity (ANOSIM) and permutational analysis of variance (PERMANOVA) were used to estimate differences between communities, using weighted UniFrac distance to partition community variability across spatiotemporal categories, i.e., river, reach, season, and their combinations. Distance-based redundancy analysis (dbRDA) was performed to investigate whether environmental variables influence changes in the biodegradation profile and multi-trophic community composition.

We used multiple co-inertia analysis (mCIA) with the omicade4 v.1.38 package [42] to estimate the relationship (global concordance) between multi-trophic communities and the biodegradation profile. Additionally, linear regression analysis, Procrustes, and Mantel tests were performed to assess the relationships between the biodegradation profile and trophic groups. Mediation linkage was evaluated using the multivariate omnibus distance mediation analysis (MODIMA) method [43] to estimate the proportional influence of benthic microbial communities on the direct effects of environmental variables on the biodegradation profiles of 96 organic compounds. A more detailed description of the downstream statistical analyses is provided in **Supplementary Text S4**.

## Results

### The biodegradation profile of organic pollutants varied across rivers, reaches, and seasons

Tian et al. [31] assessed the biodegradation potential of two rivers and their reaches (i.e., upstream and downstream segments) across four seasons by measuring the biodegradation rate constants of 96 organic compounds through batch incubation experiments. The compounds were classified into structure-based chemical taxonomic categories (e.g., superclasses) [34] to systematically organize chemical diversity and enable comparison among structurally related pollutants. These classifications summarized mixture composition and provided a framework for interpreting biodegradation variability, as shared traits (e.g., aromaticity, heteroatoms, or acidity) influence microbial accessibility, enzymatic pathways, and bioavailability. At the superclass level, benzenoid compounds had the most representation in the pollutant mixture (60.4%), followed by organoheterocyclic compounds (19.8%) and organic acids and their derivatives (9.4%) (**Figure 1B**). Other superclass categories were represented with <5.2%. We used differential analysis to identify in which River × Reach × Season group each compound’s biodegradation rate differed (**Figure 1C** and **Supplementary Table S8**). Certain superclasses were degraded across the groups, i.e., benzenoids, organic acids and derivatives, organoheterocyclic compounds, lipids and lipid-like molecules, and organic nitrogen compounds, however, their biodegradation rates varied (**Figure 1D**). At the class level, compounds under benzene and their substituted derivatives were the most prominent across the groups (**Supplementary Figure S7**).

### Spatial variation explained more variance in the benthic microbial community composition than seasonal differences in the studied rivers and reaches

In total, we identified 2,661 prokaryotic and 2,774 eukaryotic species (including ASVs assigned as unclassified taxa) from the 16S rRNA and 18S rRNA amplicon sequencing data (**Supplementary Tables S5** and **S6**). For the prokaryotes, 2.7% of the species are archaea, with 97.3% assigned as bacteria. For the eukaryotes, 27.9% of the species are algae, 26.6% are fungi, 34.6% are protists, and 10.8% are metazoans. The prokaryotes were classified into 87 phyla, with Pseudomonadota as the dominating phylum, followed by Actinomycetota and Chloroflexota (**Supplementary Figure 8A**). The eukaryotes were classified into 30 divisions, with the metazoan division, Opisthokonta, dominating most of the eukaryotic reads, followed by three protistan representatives from the TSAR supergroup, i.e., Stramenopiles, Alveolatea, and Rhizaria (**Supplementary Figure 8B** and **S9**). Both communities exhibited variable relative abundance distributions across the river, reach, and seasonal groups, with a more pronounced difference in the eukaryotic communities for the Opisthokonta and Stramenopiles supergroups (**Supplementary Figure 8B**).

We estimated diversity and community composition indices to examine the spatiotemporal variations in the benthic microbiome across trophic groups. Alpha diversity metrics (**Supplementary Table S9** and **Figure S10**) showed negative correlations between prokaryotic and eukaryotic communities, though only richness was significant (*R* = −0.3850, *p* = 0.01; **Figure 2A**). Prokaryotic alpha diversity was lowest in autumn samples, but no consistent seasonal pattern emerged, suggesting spatial variation was more influential than seasonality. Principal coordinate analysis (PCoA) of weighted UniFrac distances revealed strong differences in phylogenetic composition among River × Reach × Season groups (**Figure 2B-F** and **Supplementary Table S10**). Prokaryotes showed the greatest dissimilarity (ANOSIM *R* = 0.89, *p* = 0.001), followed by fungi (*R* = 0.85), algae (*R* = 0.80), protists (*R* = 0.72), and metazoans (*R* = 0.60).

**Figure 2.**
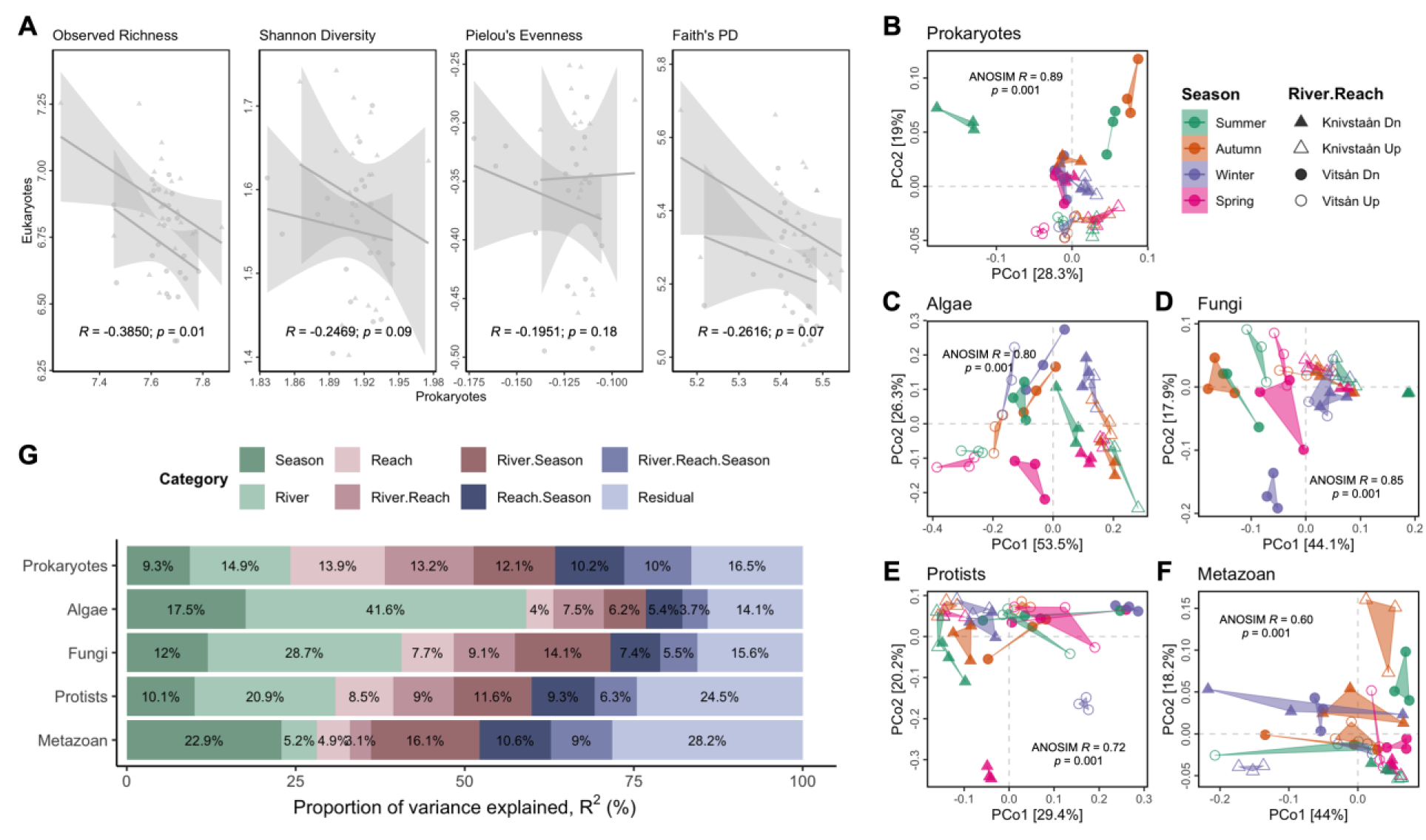
Spatiotemporal patterns of the benthic microbial communities across the rivers, reaches, and seasons. (A) Linear regression relationships between the alpha diversity metrics of the prokaryotic and eukaryotic communities. The regression line is visualized by the river category. (B-F) Principal coordinate analysis (PCoA) based on weighted UniFrac distance and analysis of similarities (ANOSIM) of each trophic community at the spatiotemporal River × Reach × Season grouping. (G) Sources of overall variation of the multi-trophic communities across spatiotemporal categories. Proportion of variance based on permutational analysis of variance (PERMANOVA) tests on weighted UniFrac distance.

Furthermore, we estimated the sources of overall variation to assess the proportional influence of the spatiotemporal variables on each trophic community using a permutational analysis of variance (PERMANOVA) on weighted UniFrac distance (**Figure 2G** and **Supplementary Table S10**). The influence of spatial variables, i.e., river (15%, PERMANOVA *R^2^* = 0.1491, *Df* = 1, *F* = 28.97, *p* = 0.001), reach (14%), and their combination (River × Reach, 13%) was relatively stronger than season (9%) on the composition of prokaryotes. Likewise, algae, fungi, and protists had a relatively higher proportional influence from rivers on their composition, with 42%, 29%, and 21%, respectively. Only the metazoan communities exhibited the highest seasonal influence, accounting for 23% of the total variation. Together, the diversity metrics and community grouping analyses indicate a consistent pattern in which spatial factors (river and reach) account for greater variation than seasonality across prokaryotic and eukaryotic trophic groups.

### Various environmental factors influence the biodegradation profile and microbial community dynamics

Given that spatiotemporal factors are also defined by differences in environmental variables (**Supplementary Figure S1**), we examined the relationships between environmental and water chemistry parameters and the composition of each trophic group. The parameters were assembled from co-located field measurements (i.e., pH, dissolved oxygen, electrical conductivity EC, and water temperature) and proxy datasets (HYPE and national monitoring) as detailed in **Supplementary Text S1**. We performed a redundancy analysis (RDA) to illustrate the relationship between environmental variables and the biodegradation rates of the compounds at the superclass level (**Figure 3A**). All variables significantly affected the biodegradation profile, except flow rate (*Pearson’s r* = 0.0795, *adj. p-value* = 0.067). Moreover, a distance-based redundancy analysis (dbRDA) based on Bray-Curtis dissimilarity revealed that 73% of the variance in the biodegradation profile was significantly influenced by pH (dbRDA, *R^2^* = 0.7302, *p* = 0.001), followed by electrical conductivity (*R^2^* = 0.4193, *p* = 0.001), chloride (*R^2^* = 0.41, *p* = 0.001), and ammonium (*R^2^* = 0.4043, *p* = 0.001), with flow rate, dissolved oxygen, total suspended solids, total nitrogen, total phosphorus, the sum of nitrate-nitrogen and nitrite-nitrogen, lead, chromium, nickel, and zinc having significant influence (p > 0.05) (**Figure 3B, Supplementary Figure S11** and **Table S11**).

**Figure 3.**
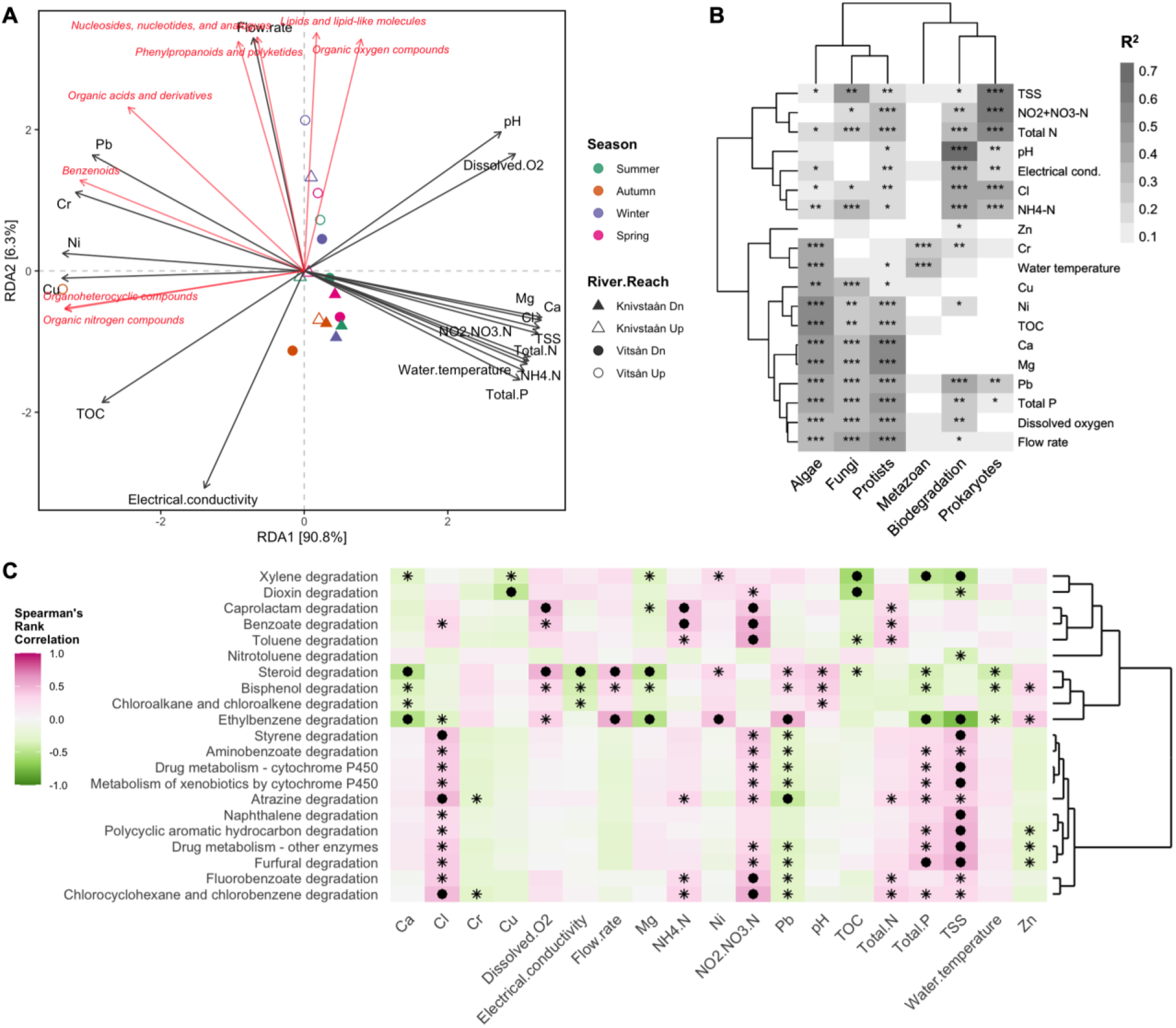
Environmental associations of the biodegradation profile and the benthic microbial communities. (A) Redundancy analysis (RDA) of the relationship between the environmental variables and the biodegradation rates (k_TCC,pH7_) of the compounds at the superclass category. (B) Proportion of the effect sizes (R^2^) of significant covariates on community profile variation from a distance-based redundancy analysis (dbRDA). The biodegradation profile was based on Bray-Curtis dissimilarity, and the trophic communities were based on weighted UniFrac distance. Hierarchical clustering using Ward’s method and Euclidean distance matrix. (C) Spearman’s rank correlation showing the relationship between the environmental variables and the predicted functions based on KEGG metabolic pathways (annotations under Level 2 – Xenobiotics biodegradation and metabolism). The asterisk indicates *P* < 0.05, and the filled circle indicates *FDR-corrected P* < 0.05. Environmental factors were log10-transformed (except pH): pH, dissolved oxygen (DO, mg/L), electrical conductivity (EC, μS/cm), and water temperature (°C) (Tian et al., 2024). Water chemistry parameters extracted from the national monitoring database: flow rate (L/s), calcium (Ca, mg/L), chloride (Cl, mg/L), chromium (Cr, µg/L), copper (Cu, µg/L), magnesium (Mg, mg/L), ammonium nitrogen (NH₄-N, µg/L N), nitrate-nitrite nitrogen (NO₂+NO₃-N, µg/L), nickel (Ni, µg/L), lead (Pb, µg/L), total suspended solids (TSS, mg/L), total organic carbon (TOC, mg/L), total nitrogen (Total N, µg/L), total phosphorus (Total P, µg/L), and zinc (Zn, µg/L).

Further hierarchical clustering analysis revealed that the biodegradation profiles clustered with the prokaryotes, due to the significant influence of similar covariates on variation in community profiles. On the other hand, the algal, fungal, and protist communities clustered together, sharing most of the environmental variables that significantly influenced their community composition, except for certain parameters, e.g., pH (associated with protists) and chromium (associated with algae). These observations indicate a relationship between environmental factors and the structure of multi-trophic benthic communities, which, in turn, shape the biodegradation of complex organic mixtures. The clustering patterns among trophic groups further suggest that shared environmental drivers govern community assembly and functional outcomes. Although the use of modeled proxy data constrains the precision of environmental associations, the analyses remain valuable for identifying consistent ecological patterns and broad environment–biodegradation linkages.

Functional profiles were predicted from KEGG pathways (**Supplementary Table S7**), yielding 383 level-3 pathways, including 21 involved in xenobiotic biodegradation (KEGG 09111). The top functions were benzoate, aminobenzoate, chloroalkane/chloroalkene, and caprolactam degradation, as well as drug metabolism (**Supplementary Figure S12**). Among these, ethylbenzene degradation showed the strongest correlations with environmental variables, e.g., total suspended solids (TSS) and total phosphorus (FDR-corrected p < 0.05; **Figure 3C**). We also assessed the beta diversity of the predicted functional profile based on Bray-Curtis dissimilarity, and found a significant influence of the spatiotemporal factors on the distribution of metabolic functions across different groups (ANOSIM *R* = 0.70, *p* = 0.001), with the River × Reach factor (35%, PERMANOVA *R^2^* = 0.3453, *Df* = 1, *F* = 87.08, *p* = 0.001) showing the highest proportional influence on its composition (**Supplementary Figure S13** and **Table S10**).

### Assessing the relationships between the biodegradation profile and multi-trophic communities

We found that the Shannon diversity of the biodegradation profile was significantly positively correlated with prokaryotes (*R* = 0.3914, *P* = 0.01) and negatively correlated with fungal communities (*R* = −0.3023, *P* = 0.04) (**Figure 4A**). No significant relationships were observed for the other communities. On the other hand, the biodegradation profile was significantly correlated (*P* < 0.001) with the composition of all multi-trophic communities, based on the Bray-Curtis index, and the functional profile, based on the Jaccard distance, with a relatively low proportion of variance explained (*R^2^* > 0.23) (**Supplementary Figure S14**).

**Figure 4.**
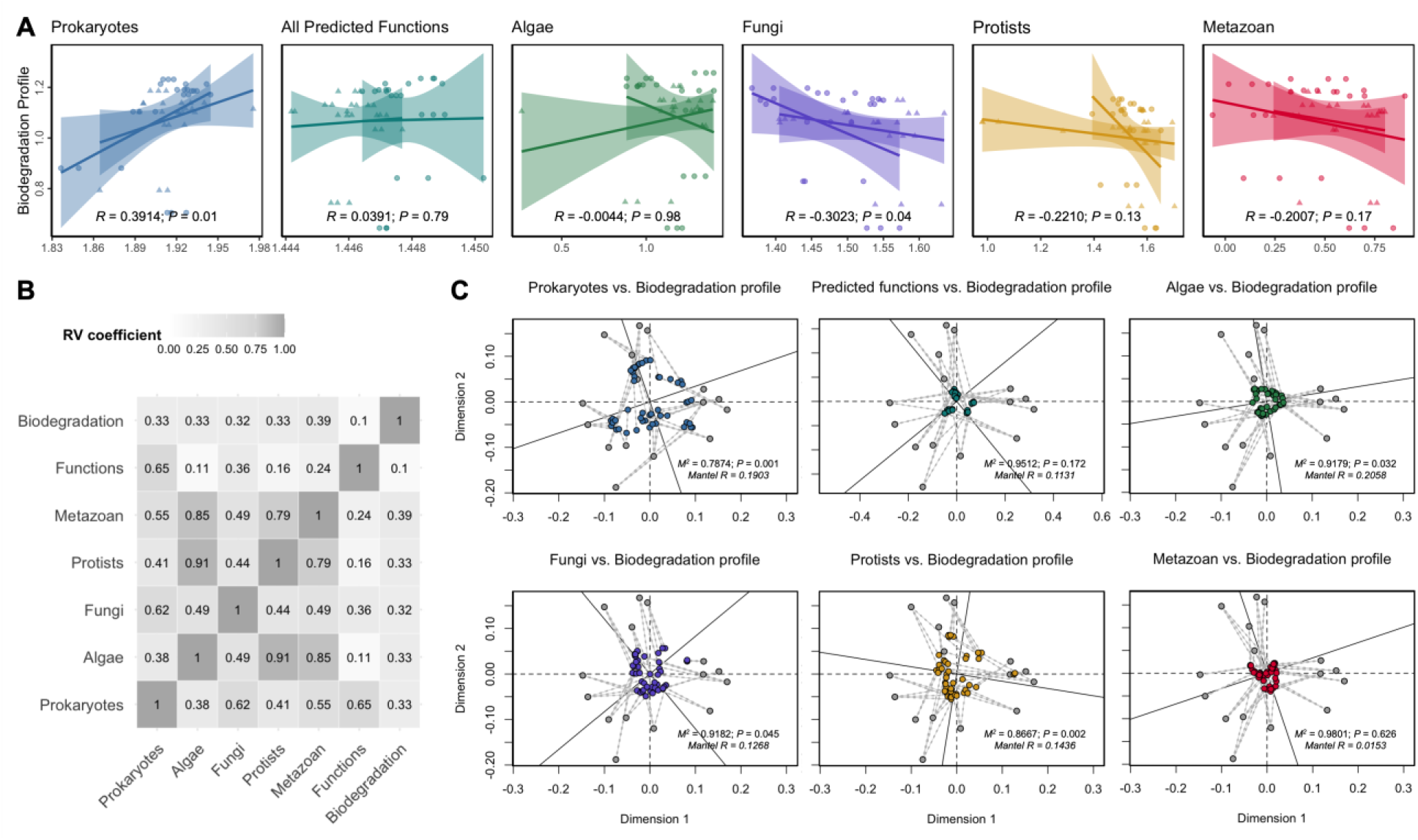
Correlations between the biodegradation profile and benthic microbial communities. (A) Linear regression relationships based on the Shannon diversity index of the biodegradation profile against each trophic community and the predicted metabolic functions. The regression line is visualized by the river category. (B) Correlations between the datasets identified by multiple co-inertia analysis (mCIA). The RV coefficient ranges from 0 to 1, with zero indicating no similarity between datasets. (C) Procrustes analysis showing the correlations between the biodegradation profile and each trophic community and predicted metabolic functions. The multi-trophic communities are based on weighted UniFrac, predicted functions using Jaccard, and biodegradation profiles using Bray-Curtis dissimilarity distance.

To further identify common relationships and assess concordance among the datasets, we performed multiple co-inertia analysis (mCIA). The RV coefficient, a measure of global similarity, was calculated to characterize the overall correlation between the biodegradation profile and multi-trophic communities (**Figure 4B** and **Supplementary Figure S15**). Notably, all multi-trophic communities showed values around 0.3-0.4, indicating low similarity, which is consistent with the beta diversity correlations. Accordingly, Procrustes analysis revealed weak clustering patterns between the biodegradation profile and each trophic group (**Figure 4C** and **Supplementary Table S13**), although they exhibited a significant clustering pattern, with prokaryotes having the lowest *M^2^* value of 0.78 (*P* = 0.001, and Mantel *R* = 0.1903), followed by protists, algae, and fungi, that had significant *M^2^* values ranging from 0.8 to 0.9. The metazoan communities and the predicted functions had no significant clustering pattern with the biodegradation profile.

### Multi-trophic communities mediate the influence of environmental factors on the biodegradation profile of rivers

We conducted mediation analyses to estimate the effect of environmental factors (independent variable) on the biodegradation profile (dependent variable), with benthic microbial communities serving as the mediators (**Supplementary Table S14**). The prokaryotic communities mediate the influence of the environmental factors on the biodegradation profile of the complex mixture of 96 organic compounds by 60% (*P_medi_* = 0.0001) (**Figure 6A**). We also identified 163 prokaryotic ASVs that were differentially associated with a River × Reach × Season group (**Supplementary Table S14**) and showed a similar percentage of mediation effect (60%, *P_medi_* = 0.0001), indicating that the differentially abundant prokaryotes included in each group had a mediation influence similar to that of the entire community. Additionally, with the predicted metabolic functional data based on KEGG pathways as mediators accounted for 30.3% (All predicted functions, *P_medi_* = 0.0001) and 29.9% (xenobiotics biodegradation and metabolism, *P_medi_* = 0.0001) of the total effect of the environment on the biodegradation profile (**Figure 6A**). Notably, the eukaryotic taxa also had significant indirect mediation effects. The algal communities accounted for 26% (*P_medi_* = 0.0001), fungi for 56% (*P_medi_* = 0.0001), protists for 53% (*P_medi_* = 0.0001), and metazoans for 38% (*P_medi_* = 0.0001) (**Supplementary Figure S16**). We also used mediation analysis to assess multi-trophic interactions and estimate how eukaryotic taxa influence the effects of environmental factors on prokaryotic communities (**Figure 6B**). Fungal and protist communities showed strong mediation effects (83% and 73%, *P* < 0.0001), indicating a major role in shaping prokaryotic assembly. Algae and benthic metazoans had weaker but still significant effects (39% and 25%), suggesting more modest contributions.

We further assessed the relationships between the differentially abundant prokaryotes associated with the spatiotemporal categories (**Supplementary Figure S17** and **Table S14)** and the biodegradation profile of the compounds. The association between the indicator species and each compound is presented in **Supplementary Figure S18.** To better visualize this association, we assessed the relationship at the phylum level for prokaryotes and at the superclass level for the compounds (**Figure 6A**). Notably, the biodegradation rates of benzenoids, nucleosides, nucleotides, and analogues, as well as organic acids and derivatives, showed strongly significant positive correlations (*FDR-corrected P* < 0.05) with Nitrospirota, Nitrospinota, RCP2-54, MBNT15, Spirochaetota, and Halobacteriota. Pseudomonadota and Verrucomicrobiota were positively correlated with the degradation of lipids and lipid-like molecules, while Chloroflexota was positively correlated with the degradation of organic nitrogen and organoheterocyclic compounds. For the predicted functions related to xenobiotic biodegradation (**Figure 6B**), the biodegradation rates of benzenoids are positively correlated with functions involved in the degradation of chloroalkanes and chloroalkenes, while also negatively associated with the degradation of chlorocyclohexanes and chlorobenzenes.

## Discussion

Recent studies have demonstrated the impact of spatial variation and temporal shifts on pollutant transformation in aquatic systems [e.g., 31, 44, 45], but the role of the biological communities involved in these processes remains understudied. The complex interaction among microbial community dynamics, environmental factors, and complex pollutant mixtures governs the resilience and functioning of river ecosystems [46, 47]. Spatiotemporal variations in community composition, driven by fluctuations in water chemistry, nutrient inputs, and hydrological dynamics, may affect microbially mediated degradation pathways and overall ecosystem multifunctionality [17, 26]. Therefore, it is crucial to examine how community dynamics and spatiotemporal variation influence ecosystem resilience and the efficiency of microbially mediated biodegradation of organic pollutants.

In this study, we demonstrated that the biodegradation profile of a complex mixture of 96 organic compounds in rivers and their reaches across four seasons is not solely influenced by direct environmental impacts but is mainly mediated by benthic microbial communities that include multi-trophic levels. Using mediation analysis, we explained how the response variable (biodegradation) occurs in sequence through mediators, supporting causal rather than merely descriptive interpretations [48]. We report that prokaryotic communities accounted for approximately 60% of the total effect of key environmental drivers on biodegradation outcomes. Notably, the same contribution was observed when only a subset of 163 differentially abundant ASVs was considered (**Figure 5A** and **Supplementary Table S14**). This suggests that the indicator taxa identified were directly involved in mediating the influence of environmental factors on the biodegradation capacity of the rivers across spatiotemporal gradients. These observations align with previous reports highlighting the pivotal role of prokaryotes in pollutant transformation in aquatic systems [13, 47, 49].

**Figure 5.**
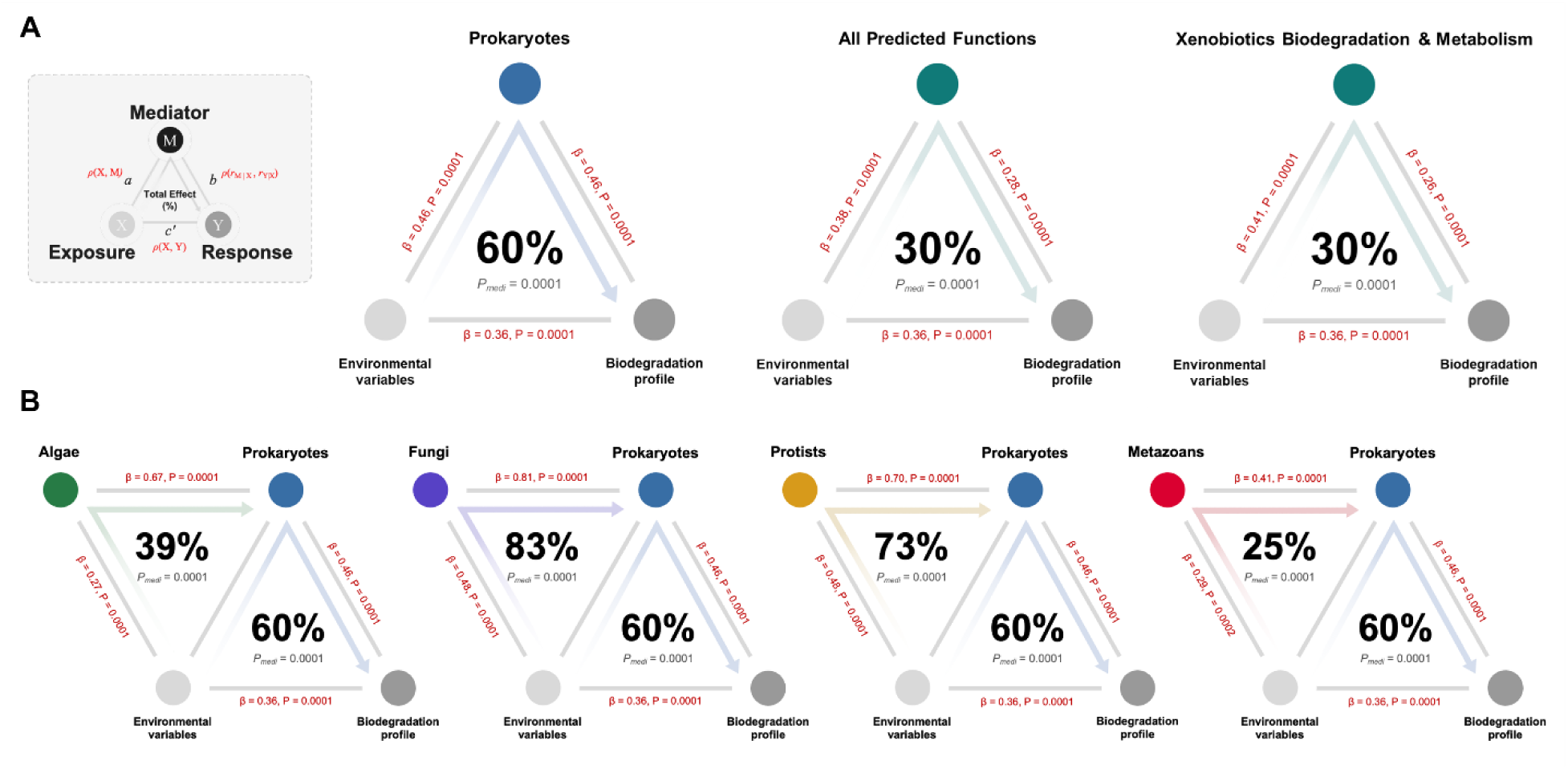
Mediation linkage between the environmental variables, biodegradation profile, and the benthic microbiome. The mediation association of the (A) prokaryotic communities and their predicted functions as mediators of the link between the environmental variables (exposure factor) and the biodegradation profile of the 96 organic compounds (response). (B) Presents the mediation association of each eukaryotic trophic group as the mediators of the influence of environmental factors on the prokaryotic communities. Beta coefficients and P-values are listed at each connecting line, and the proportions of indirect effect (mediation effect) and mediation *P*-values (*P_medi_*) are presented at the center of the ring charts.

Additionally, the predicted functional profile of prokaryotes accounted for 30.3% of the overall environmental mediation, while pathways specifically associated with xenobiotic biodegradation and metabolism accounted for up to 29.9% of the effect (**Figure 5A**). These suggest that even a modest set of xenobiotic metabolism genes can account for a significant portion of the observed degradation outcomes, demonstrating a link between functional genetic capacity within the prokaryotic fraction and pollutant biodegradation [13, 50, 51]. Key enzymatic players within these pathways, e.g., dioxygenases, dehydrogenases, and monooxygenases, catalyze critical reactions in the breakdown of organic compounds [52, 53]. Such enzymes enable the transformation of complex pollutants, ranging from aromatic hydrocarbons (e.g., benzoate, toluene, and styrene) to hazardous compounds (e.g., dioxins and chloroalkanes), into intermediates that are then channeled into central metabolic processes [54].

Beyond the direct influence of prokaryotes, our analyses extend to the indirect roles of eukaryotic taxa. Algal, fungal, protistan, and metazoan communities each showed statistically significant mediation effects on the biodegradation profile, with fungi and protists contributing most strongly (56% and 53%; **Supplementary Figure S16**). These results indicate that fungi and protists are key regulators of microbial networks that drive pollutant degradation. Fungi are known to play a significant role in riverine pollutant biodegradation of hydrocarbons, pesticides, and heavy metals through enzymatic processes and physical structures [55, 56]. Whereas protists are known to enhance pollutant biodegradation in rivers through predation [19, 57], increasing nutrient release, and promoting the activity of pollutant-degrading bacteria while also directly consuming biodegradable pollutants [58]. Additionally, algal and metazoan communities contributed more modestly (26% and 38%, respectively). Nonetheless, they have been reported to provide critical support functions for oxygen production, biofilm formation, and sediment mixing [22, 59]. Furthermore, mediation analyses also showed that fungi and protists exert particularly potent effects on prokaryotic community structure: fungal communities mediated 83% of the environmental impact on prokaryotes, while protists mediated 73%. These pronounced effects are supported by well-documented mechanisms, including the secretion of extracellular enzymes [60], nutrient recycling through the degradation of complex substrates [61], and predation that enhances bacterial turnover and nutrient release [57, 62]. Although the mediation effects attributed to algal and metazoan communities were lower (39% and 25%, respectively), their roles remain ecologically significant, contributing to bioturbation and biodeposition that maintain sediment structure [63, 64].

Most of the organic compounds in the pollutant mixture assessed in this study are categorized as benzenoids (60.4%) based on chemical structure (**Figure 1B**). Accordingly, based on our metabolic functional predictions of prokaryotic communities, degradation of benzoate, aminobenzoate, chloroalkane and chloroalkene, caprolactam, and drug metabolism (other enzymes) are the most prominent biodegradation processes. Previous studies have found benzoate degradation to be a key metabolic pathway in rivers, particularly a central pathway for the catabolism of most aromatic compounds [50, 65]. By correlating the predicted metabolic functions and the biodegradation profile (**Figure 6B**), we found that the biodegradation rates of benzenoids and the functions responsible for degrading chloroalkanes and chloroalkenes were positively correlated. In contrast, functions that target chlorocyclohexanes and chlorobenzenes were negatively correlated. Similarly, the degradation of organoheterocyclic compounds was significantly associated with the expression of cytochrome P450 enzymes as well as pathways involved in aminobenzoate and styrene degradation. These associations not only provide mechanistic insights into how distinct chemical features are targeted by microbial enzymatic systems but also highlight a high degree of substrate specificity in biodegradation [51].

**Figure 6:**
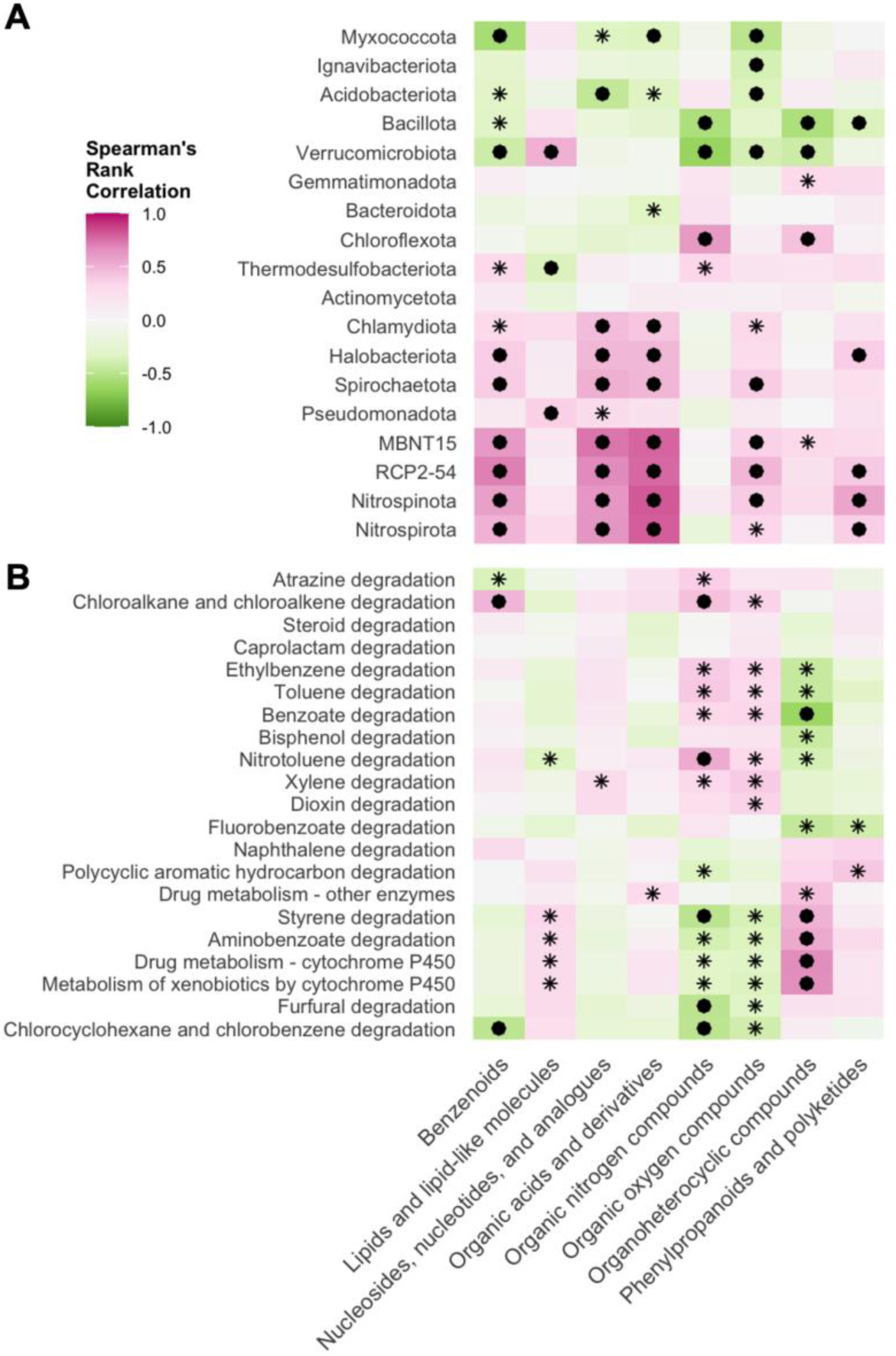
Relationships between indicator prokaryotic taxa and their predicted functions and the biodegradation profile. Spearman’s rank correlation showing the relationship between the biodegradation rates of the organic compounds at the superclass level and the (A) prokaryotic phyla of the spatiotemporally abundant taxa across the River × Reach × Season group, and (B) the predicted functions based on KEGG metabolic pathways (annotations under Level 2: Xenobiotics biodegradation and metabolism). The asterisk indicates *P* < 0.05, and the filled circle indicates *FDR-corrected P* < 0.05.

Exploring the taxonomic factors influencing biodegradation, we found that the biodegradation rates of specific compound classes are strongly correlated with the relative abundances of distinct prokaryotic phyla (**Figure 6A**). For instance, biodegradation of benzenoids, nucleosides, nucleotides, and related organic acids was positively associated with members of the bacterial sister phyla Nitrospirota and Nitrospinota, Spirochaetota, and Halobacteriota. These lineages are equipped with catabolic pathways optimized for the breakdown of aromatic and acidic functional groups [49, 66, 67]. Concurrently, Pseudomonadota and Verrucomicrobiota exhibited positive correlations with the degradation of lipids and lipid-like molecules, supporting the notion that these groups contribute to the hydrolysis and further oxidation of hydrophobic pollutants [49, 68]. Furthermore, Chloroflexota showed significant positive associations with the degradation of organic nitrogen compounds and organoheterocyclic molecules, suggesting specialization in targeting nitrogenous or heterocyclic moieties that are typically more recalcitrant [69]. Nevertheless, we acknowledge the limitations of predicting microbial functional potential and activity from prokaryotic taxonomy. Taxonomic identity may not fully reflect the samples’ functional capacity because it does not account for horizontal gene transfer, metabolic plasticity, or strain-level variation. Therefore, we recommend further combining metagenomic and metaproteomic methods to directly evaluate microbial functional potential and activity.

We also observed that spatial variation had a greater impact on benthic microbial communities across multi-trophic groups than seasonality. This underscores that spatial heterogeneity, driven by riverine and reach-specific environmental factors, plays a pivotal role in shaping benthic microbial diversity and composition, thereby establishing distinct biogeographical patterns in river ecosystems [70, 71]. This also reinforces the concept that spatially structured environmental gradients are the primary drivers of benthic microbial community assembly, whereas temporal fluctuations exert only secondary effects [72, 73]. Hence, while temporal variation is important, spatial processes, e.g., habitat heterogeneity, dispersal limitation, and species sorting, are significant drivers of river microbial assembly, often overriding temporal changes [74]. However, we note that our spatial inference was limited to two rivers and two reaches, and because river-specific attributes may inflate these spatial effects, we recommend a broader spatial replication to further generalize these findings.

Our study reveals profound ecological implications, as both prokaryotic and eukaryotic communities collectively mediate the majority of the environmental impact on biodegradation. Consequently, any perturbation, whether arising from spatial variation, altered nutrient loading, temperature fluctuations, or increased contaminant inputs, can have cascading effects throughout the microbial food web [75]. This indicates that river management should focus not only on pollution control but also on restoring ecosystem health (e.g., through bioremediation) using an ecosystem-based approach [76, 77]. Additionally, the strong mediation effects observed for fungi and protists suggest that enhancing bioremediation outcomes may be achieved by targeting these groups for conservation or bioaugmentation. For example, approaches that stimulate natural fungal growth or promote protistan activity may yield more robust and resilient prokaryotic consortia capable of efficient pollutant degradation [78]. Moreover, the detailed associations between specific prokaryotic taxa and the degradation of distinct chemical classes carry practical implications for monitoring and managing contaminated river systems [9, 13]. By using taxa, e.g., Nitrospirota or Pseudomonadota as bioindicators, it may be possible to develop systems that signal shifts in biodegradation potential before toxic pollutant concentrations accumulate [79]. Likewise, understanding the molecular targets, as revealed by the metabolic pathway associations, provides opportunities for targeted bioaugmentation strategies that can selectively enhance the degradation of particularly recalcitrant pollutant fractions [80]. Moreover, our results suggest that management strategies designed to enhance the natural attenuation capacity of aquatic systems should consider the complex interactions between environmental drivers and microbial community assembly. In urban and industrial environments, where rivers are often burdened with organic pollutants from wastewater and stormwater, it is especially important to understand how microbes mediate these processes. Initiatives aimed at restoring polluted rivers to a more natural state should therefore consider the entire microbial food web, with particular attention paid to both the prokaryotic degraders and their eukaryotic modulators [81].

Moving forward, the multilayered mediation effects identified in our study underscore the need for further research that integrates multi-omic tools. Integrative studies would enable a more comprehensive understanding of the molecular pathways and regulatory networks that underlie biodegradation in complex ecological settings [82]. Additionally, we used 18S rRNA genes, which offered limited taxonomic resolution for fungi compared to the Internal Transcribed Spacer (ITS) region. To enhance future research, we recommend incorporating ITS-based profiling alongside 18S rRNA gene profiling to improve fungal identification and ecological interpretation. Coupling molecular techniques with detailed assessments of environmental parameters and microbial community structure could lead to the development of predictive models that not only forecast biodegradation outcomes but also inform the design of targeted bioremediation interventions. Addressing these limitations will improve both the resolution and reliability of multi-trophic ecological insights.

In conclusion, our findings advance our understanding of the ecological mechanisms underlying microbially mediated pollutant degradation and underscore the need for integrated management strategies that account for the entire microbial food web. Such an approach will be essential for the effective management and restoration of polluted river ecosystems in the face of ongoing anthropogenic and climate-driven changes.

## Acknowledgments

We thank Michael S. McLachlan for his valuable insights on the manuscript’s initial draft. The computations were performed on the supercomputer Dardel at PDC in KTH Royal Institute of Technology, with access and resources provided by the National Academic Infrastructure for Super-computing in Sweden (NAISS) through projects NAISS 2025/22-1105 and 2025/23-467.

## Author contributions

J.M.S., E.B., B.D., and M.P. conceptualized the study. M.P. and R.T. conceptualized the sampling and experimental design and conducted field sampling. R.T. performed biodegradation experiments, processed, and analyzed the chemical data. J.M.S. performed DNA preparation for sequencing, analysis of amplicon sequence data, downstream analyses, and wrote the initial manuscript draft. B.D. retrieved and pre-processed the physico-chemical and hydrological data. All authors read, revised, and approved the final version of the manuscript.

## Data availability

Raw sequencing data are deposited at the NCBI SRA archive under project PRJNA1305693. The corresponding visualization and analysis data, as well as the R codes, are publicly accessible at https://github.com/jserrana/spatiotemporal-bie.

## Consent for publication

Not applicable.

