## Supplementary Information for "Pollutant biodegradation profile mediated by multi-trophic microbial dynamics in rivers"

Running Title: Spatiotemporal biodegradation in rivers

#### Authors & Affiliations

Joeselle M. Serrana[1,2]*, Run Tian[2,3], Francisco J. A. Nascimento[1,4,5], Elias Broman[1,4,5], Benoît Dessirier[1,5], and Malte Posselt[1,2]

[1] Stockholm University Center for Circular and Sustainable Systems (SUCCeSS), Stockholm University, 106 91 Stockholm, Sweden

[2] Department of Environmental Science, Stockholm University, 106 91 Stockholm, Sweden

[3] Queensland Alliance for Environmental Health Science (QAEHS), The University of Queensland, 20 Cornwall Street, Woolloongabba, Queensland 4102, Australia

[4] Department of Ecology, Environment, and Plant Sciences, Stockholm University, 106 91 Stockholm, Sweden

[5] Baltic Sea Centre, Stockholm University, Stockholm, Sweden

#### Supplementary Table Legends

**Supplementary Table S1.** Sample description based on spatial and temporal category, and their environmental variables: pH, dissolved oxygen (DO, mg/L), electrical conductivity (EC, µS/cm), and water temperature (°C) (Tian et al., 2024). Water chemistry parameters extracted from the national monitoring database: flow rate (L/s), calcium (Ca, mg/L), chloride (Cl, mg/L), chromium (Cr, µg/L), copper (Cu, µg/L), magnesium (Mg, mg/L), ammonium nitrogen ($NH_4$-N, µg/L N), nitrate-nitrite nitrogen ($NO_2$+$NO_3$-N, µg/L), nickel (Ni, µg/L), lead (Pb, µg/L), total suspended solids (TSS, mg/L), total organic carbon (TOC, mg/L), total nitrogen (Total N, µg/L), total phosphorus (Total P, µg/L), and zinc (Zn, µg/L).

**Supplementary Table S2.** Biodegradation rate constants from Tian et al. (2024). Compound classification based on structure-based chemical taxonomic categories.

**Supplementary Table S3.** Metabarcoding information for the 16S and 18S rRNA amplicon sequencing datasets.

**Supplementary Table S4.** Eukaryotic grouping categories for the 18S rRNA dataset to assign ASVs to algae, fungi, protists, and metazoan taxa.

**Supplementary Table S5.** 16S rRNA amplicon sequencing data: ASV table and taxonomic assignment.

**Supplementary Table S6.** 18S rRNA amplicon sequencing data: ASV table and taxonomic assignment.

**Supplementary Table S7.** Predicted functions based on the KEGG pathway database.

**Supplementary Table S8.** Differential abundance analysis of the biodegradation profile.

**Supplementary Table S9.** Alpha diversity estimates of the multi-trophic groups.

**Supplementary Table S10.** Permutational analysis of variance (PERMANOVA) and analysis of similarities (ANOSIM) statistics of the multi-trophic groups based on the River × Reach × Season category.

**Supplementary Table S11.** Distance-based redundancy analysis (dbRDA) statistics of the multi-trophic groups based on the River × Reach × Season category.

**Supplementary Table S12.** Multiple co-inertia analysis (mCIA) statistics of the multi-trophic groups.

**Supplementary Table S13.** Procrustes and Mantel test statistics of the multi-trophic groups.

**Supplementary Table S14.** Mediation analysis statistics based on multivariate omnibus distance mediation analysis (MODIMA).

**Supplementary Table S15.** Differential abundance analysis by Random Forests to identify indicator taxa based on the River × Reach × Season category.

### Supplementary Text

**Supplementary Text S1.** Environmental variables

The environmental variables recorded in the field were pH, dissolved oxygen (DO, mg/L), electrical conductivity (EC, μS/cm), and water temperature (°C) (**Supplementary Table S1** and **Figure S1**). For detailed site and sampling information, see Tian et al. (2024). Additional hydrological and chemical data were added from national databases. Daily modeled streamflow values were extracted from a national model shared publicly by the Swedish Meteorological and Hydrological Institute (SMHI) (https://vattenwebb.smhi.se/nadia/); for this study, we used data produced by the model HYPE_version_5_19_0, and the model setup s-hype2016_version_16_i, for the areas with SUBID 40831 (Vitsån, outlet) and SUBID 9026 (Knivstaån, just downstream of Knivstaån_Dn). Fourteen water chemistry parameters were extracted from the Swedish environmental monitoring database (Miljödata-MVM, 2024) for stations *Knivstaån uppströms reningsverk* (MVM id 45499), *Reningsverket syd* (MVM id 45491), *Rocklösaån R3* (MVM id 44872), and *Hågaån, nedströms ARV* (MVM id 44498).

The locations for the modeled streamflow and the national monitoring of chemical parameters do not coincide perfectly. For Knivstaån, the two national monitoring stations correspond almost directly to the upstream (Knivstaån_Up) and downstream reach (Knivstaån_Dn) of Knivstaån, respectively, with the reference point for streamflow fairly close to Knivstaån_Dn. Knivstaån_Up and Knivstaån_Dn are only 600 m apart, with no stream confluence between the two, which makes it so that catchment areas draining to the two reaches are very similar, and we could consider that the same modeled streamflow values were representative of both reaches. Furthermore, the parameter values from the monitoring were available by season for at least three consecutive years 2017-2019, sometimes for 2017-2022, which informed not only about the expected seasonal levels but also about their possible interannual variability. For Vitsån, the situation is more complex: two branches, Rocklösaån and Hågaån, merge at a confluence to form the river Vitsån. The reference point for the modeled streamflow is located approximately 5 km downstream of the confluence. A catchment delineation was calculated for each branch of Vitsån and the outlet using GRASS-8.3 functions *r.terraflow* (flow accumulation) and *r.water.outlet* (catchment delineation for each reach) on the 2m-resolution digital elevation model (2019 version) from the Swedish mapping, cadastral, and land registration authority (Lantmäteriet). This revealed that 56% of the catchment drains into Rocklösaån, 24% into Hågaån, and 20% into Vitsån downstream of the confluence. Without any systematic differentiation of the three areas in terms of geology or land use, the modeled streamflow was assumed to emanate from each sub-catchment proportionally to their surface area. In terms of stream water chemistry, Vitsån_Up is located 1.5 km upstream of *Rocklösaån R3*, with no tributaries or major effluent outlets between the two. Therefore, we directly used values at *Rocklösaån R3* to represent Vitsån_Up. Vitsån_Dn is located just downstream of the confluence of Rocklösaån (where Vitsån_Up is located) and Hågaån (where the local wastewater treatment plant actually releases its effluents and *Hågaån, nedströms ARV* is located). The representative values for environmental parameters at Vitsån_Dn were thus calculated as the weighted sum (by catchment area draining to each branch of the stream) of stations Rocklösaån (70%) and Hågaån (30%). The data availability at Vitsån was also more limited in time, with a full seasonal monitoring time-series only for year 2017; the representativity of this particular year was assessed by comparison to the monitoring time series in Husbyån, the next small river over to the North-East of Vitsån, that has seasonal time series of environmental monitoring spanning 2012-2024. These comparisons supported the concentration values from the sites in Vitsån available in the national monitoring database as suitable proxy environmental data to include in the spatio-temporal study of biodegradation.

These data, i.e., flow rate (L/s), turbidity (mg/L), total organic carbon (TOC, mg/L), nutrients, i.e., ammonium nitrogen ($NH_4$-N, µg/L N), nitrate-nitrite nitrogen ($NO_2$+$NO_3$-N, µg/L), total nitrogen (Total N, µg/L), total phosphorus (Total P, µg/L), major ions, i.e. calcium (Ca, mg/L), magnesium (Mg, mg/L), and chloride (Cl, mg/L), and metals, i.e., chromium (Cr, µg/L), copper (Cu, µg/L), nickel (Ni, µg/L), lead (Pb, µg/L), and zinc (Zn, µg/L) are provided in **Supplementary Table S1**.

Based on the modeled streamflow and stream water temperature for the period 1991-2023, a local definition of the seasons was established. Spring, starting on April 1$^{st}$, sees the water temperature rise and the streamflow decrease. The summer, starting June 10th, consists mainly of a high plateau of stream temperature and low streamflow. Autumn, starting August 20$^{th}$, exhibits a concurrent decrease in temperature and increase in streamflow. Winter, starting December 1$^{st}$, was characterized by low temperatures and high streamflow values. This local definition of seasons was used to aggregate the hydrological and chemical parameters for each reach. Following this definition, field sampling occurred near the end of each season, except for autumn, for which it took place in the middle of the season. The Pearson correlation p-values reveal statistically significant relationships among the environmental variables (**Supplementary Figure S2**). For example, water temperature had significant positive correlations with electrical conductivity, magnesium, calcium, chloride, total phosphorus, turbidity, and nitrate-nitrite nitrogen, and significant negative correlations with flow rate, pH, dissolved oxygen, total organic carbon, copper, chromium, lead, nickel, and zinc. These results highlight key interactions among physicochemical parameters, i.e., pH, dissolved oxygen, electrical conductivity, and flow rate.

**Supplementary Text S2.** Amplicon sequencing and bioinformatics

Total genomic DNA was extracted from ~300 mg of sediment from each sample using the DNEasy PowerSoil Pro kit (Qiagen GmbH, Hilden, Germany), following the manufacturer's instructions with minor modifications to the cell lysis time and elution steps. The quantity and quality of extracted DNA were measured using the NanoPhotometer N60 (IMPLEN GmbH, Germany). Aliquots of each DNA sample were then sent to Novogene Europe (Cambridge, UK) for amplicon sequencing using two barcode regions. The prokaryotic V4-V5 region of the 16S rRNA gene was amplified using the 515F-907R primers, and the eukaryotic V4 region of the 18S rRNA gene was amplified using the 528F-706R primers. All PCR reactions consisted of 10 ng of DNA template, two µM of forward and reverse primers, and 15 µL Phusion® High-Fidelity PCR Master Mix (New England Biolabs), amplified under the following conditions: initial denaturation for 1 min at 98°C, followed by 30 cycles of denaturation at 98°C for 10 s, annealing at 50°C for 30 s, elongation at 72°C for 30 s, and a final extension step of 5 min at 72°C. Quality-checked PCR products were pooled at equal densities and purified using a Qiagen Gel Extraction

Kit (Qiagen, Germany). Following the manufacturer's recommendations, sequencing libraries were generated using the TruSeq DNA PCR-Free Sample Preparation Kit (Illumina, USA), and index codes were added accordingly. The quality of the resulting amplicon library was assessed on the Qubit 2.0 Fluorometer (Thermo Scientific) and the Agilent Bioanalyzer 2100 system. The qualified amplicon libraries were then sequenced on an Illumina NovaSeq 6000 platform (Illumina, San Diego, CA, USA), generating 250 bp paired-end reads.

The raw reads were checked for quality using FastQC v.0.11.9 (Andrews, 2010). The paired-end reads were demultiplexed into each sample based on their unique barcodes and then truncated by removing the barcode and primer sequences. The demultiplexed paired-end reads were then quality-screened and processed via the DADA2 v.1.32.0 pipeline (Callahan et al., 2016). The rarefaction curves calculated for ACE, Chao1, and observed ASV indices approached a plateau or a stable point, indicating that sequencing depth was sufficient to estimate and compare diversity metrics across samples (**Supplementary Figure S3**). Read filtering was performed using the following quality parameters: a maximum expected error of 2, and minimum length of 150 bp. Reads matching against the PhiX genome were also filtered out. Amplicon sequence variants (ASVs) were inferred by denoising the quality-filtered reads with the DADA2 error model. The reads were then dereplicated, merged, and sequences containing chimeric reads were removed (**Supplementary Table S3**). ASVs with less than 10 reads were removed. For each ASV, the taxonomic assignment was performed using the *assignTaxonomy* function in DADA2, implementing the RDP naive Bayesian classifier method (Wang et al., 2007) against the Silva v.138.2 (Quast et al., 2013) database for 16S and the Protist Ribosomal Reference (PR2) v.5.1.0 (Guillou et al., 2012) database for 18S with the minimum bootstrapping support set to default. Multiple sequence alignments of the prokaryotic and eukaryotic datasets were performed using DECIPHER v.2.26.0 (Wright, 2015). A phylogenetic tree was then constructed with FastTree v.2.1.11 (Price et al., 2009) using the GTR nucleotide substitution model.

From the 10 million 16S and 9.9 million 18S raw reads, a total of 5.6 million and 6.7 million non-singleton sequences remained after quality filtering, merging paired-end reads, and removal of chimeras and singleton reads. For the 16S data, only ASVs assigned to bacteria and archaea were retained, while ASVs assigned to chloroplasts or mitochondria were discarded. For the 18S data, ASVs annotated as bacteria, archaea, or embryophytes were filtered out. Ecological categories, i.e., ecological function (parasite, phagotroph, phototroph, and metazoa) and trophic group annotations (parasitic, phototrophic, metazoan, fungi, and consumer) were assigned to the eukaryotic data by matching its taxonomic information against the ecological functions assigned to the ASV table from the metaPR2 database (Vaulot et al., 2022). The annotations in this database are adopted from the ecological classifications based on literature used by Sommeria-Klein et al. (2021), with additional assignments to taxonomic groups as defined in Vaulot et al. (2022). The datasets were then assigned to five categories: prokaryotes (bacteria and archaea), algae, fungi, metazoans, and protists (**Supplementary Table S4**). Sample sequence abundances were standardized to the median sequencing depth. The resulting ASV tables (**Supplementary Tables S5 and S6**) and phylogenetic trees for each dataset were compiled into phyloseq objects using the phyloseq v.1.48.0 package (McMurdie & Holmes, 2013) and microtables using the microeco v.1.15.0 package (Liu et al., 2021) for the downstream analyses. Using the trans_func class of microeco, Tax4Fun2 (Wemheuer et al., 2020) was used to predict functional profiles of the prokaryotic communities from the 16S data using the KEGG database (Kanehisa & Goto, 2000; Kanehisa, 2019; Kanehisa et al., 2025) (**Supplementary Tables S7**).

**Supplementary Text S3.** Field vs batch microbial profile

Microbial communities from before (field) and after (batch) the biodegradation experiment were characterized to determine if there are differences between the microbial profiles at the start of the biodegradation experiment and after the 10-day incubation. This pre-evaluation is important for understanding how incubation conditions influence microbial dynamics compared to the field profile, providing insights into the representativeness and environmental relevance of our lab-based biodegradation experimental setup. To evaluate differences in alpha diversity between the datasets, we calculated and compared several metrics: Observed ASV richness, Chao1 richness, Shannon and Simpson diversity indices, Pielou's evenness, and Faith's phylogenetic diversity. A student t-test was performed to statistically compare the two sample types. We found no significant differences in all estimated alpha diversity indices between the field and batch samples for both 16S and 18S data (**Supplementary Figure S4**).

To compare the datasets based on composition, we performed a co-inertia analysis (CIA) (Dray, Chessel & Thioulouse, 2003) using the *mCIA()* function of the omicade4 v.1.38 package (Meng et al., 2013) to assess the relationships and trends between the field and batch datasets. CIA is a multivariate method used to measure the strength of association between two sets of variables (Culhane, Perrière & Higgins, 2003). It employs a covariance-based objective function to identify common relationships and assess concordance among multiple datasets. The analysis was performed at the ASV-level. Before the CIA, the abundance data were Hellinger-transformed to reduce the weights of overly abundant ASVs per categorical group. The RV coefficient, or the measure of global similarity between the datasets, is calculated using a Monte-Carlo test on the sum of eigenvalues. Additionally, we performed beta diversity analysis to compare the community structures of the two datasets. Beta diversity was assessed based on unweighted and weighted UniFrac distance calculated with the *trans_beta* class of the microeco package. Community dissimilarities were then determined and visualized by principal coordinate analysis (PCoA). Lastly, Mantel statistic based on Pearson's product-moment correlation was computed to assess the correlation between the UniFrac distances of the two datasets using the mantel function of the vegan v.2.6-4 package (Oksanen et al., 2013). The projections of the field vs batch datasets onto the principal components of CIA are presented in **Supplementary Figure S5A-D**. The co-inertia analysis revealed strong global concordance between the field and batch datasets, with RV coefficients of 96% for 16S and 92% for 18S data, indicating highly similar multivariate structures between the two sample types. However, comparisons of beta diversity using unweighted and weighted UniFrac distances showed contrasting results. The PCoA plot of unweighted UniFrac distance for the 16S and the 18S data (**Supplementary Figure S6A-B**) demonstrated a clustering of samples between sample type, with a Mantel statistic r of 0.52 ($p = 0.001$) and 0.42 ($p = 0.001$), respectively. On the other hand, weighted UniFrac distance for the 16S and the 18S data (**Supplementary Figure S6C-**

**D**) showed significant differences by sample type, with a Mantel statistic r of 0.33 (p = 0.001) and 0.53 (p = 0.001), respectively.

To summarize, we found that the microbial profiles of the field and batch samples for both 16S and 18S data did not differ significantly across various alpha diversity metrics, i.e., richness, evenness, and phylogenetic diversity, indicating comparable within-sample diversity. This was supported by the CIA, which showed strong global concordance, suggesting a similar multivariate structure between the two sample types. Beta diversity based on unweighted UniFrac distances indicated clustering of samples on both sample types, reflecting similarities in the differences in community composition based on phylogeny; however, an abundance-based metric, i.e., weighted UniFrac distance, showed significant differences in relative abundance of taxa between the field and batch samples. This suggests that differences in the abundance of certain species between the start of the experiment and the 10-day incubation, even though the species richness and diversity remained consistent. Although the 16S and 18S species richness and diversity were consistent with strong similarity in multivariate structure between the two sample types, the community dissimilarities were not comparable based on weighted phylogenetic distance, suggesting shifts in abundance of microbial representatives from the microbial inoculum of the biodegradation experiment to the last day of incubation. Given that the biodegradation rates were based on first-order rate constants, the field samples were used to represent the microbial community profile in the subsequent analyses.

**Supplementary Text S4.** Statistical analyses

*Diversity and community composition analysis*

Data processing, visualization, and statistical analyses were performed in R v.4.4.3 (R Core Team, 2025). Using a random forest analysis, the compounds were assigned to the River × Reach × Season category, to which they are differentially associated. All 96 compounds were found to be significant after the Kruskal-Wallis rank sum test and p-value adjustment (**Supplementary Table S8**). The relative abundance of each taxon was visualized using the mean relative abundance for samples grouped by River × Reach ×

Season. Taxonomic and phylogenetic alpha diversity indices were calculated, i.e., observed richness, Shannon diversity, Pielou’s evenness, and Faith's index of phylogenetic diversity (PD) (**Supplementary Table S9**). The seasonal effects on alpha diversity were assessed with an analysis of variance (ANOVA) test.

Beta diversity of the benthic microbiome was assessed using weighted UniFrac distance and Bray-Curtis dissimilarity indices. These were estimated using the diversity-based functions of the microeco v.1.15.0 package (Liu et al., 2021). Principal coordinate analysis (PCoA) was performed to visualize the differences in community composition between groups. An analysis of similarity (ANOSIM) and permutational analysis of variance (PERMANOVA) were used to estimate the differences between communities, utilizing weighted UniFrac distance to partition the variability of communities across spatiotemporal categories, i.e., river, reach, season, and their combinations (**Supplementary Table S10**). Distance-based redundancy analysis (dbRDA) was performed to investigate whether environmental variables influence changes in the biodegradation profile and multi-trophic community composition(**Supplementary Table S11**).

*Comparison of multi-trophic groups*

We used a multiple co-inertia analysis (mCIA) with the omicade4 v.1.38 package (Meng et al., 2013) to estimate the relationship or global concordance between the multi-trophic communities and the biodegradation profile (**Supplementary Table S12**). Additionally, linear regression analysis, Procrustes, and Mantel tests were performed to assess the relationships between the biodegradation profile and trophic groups (**Supplementary Table S13**).

*Mediation analyses*

We used a multiple co-inertia analysis (mCIA) with the omicade4 v.1.38 package (Meng et al., 2013) to estimate the relationship or global concordance between the multi-trophic communities and the biodegradation profile. Additionally, linear regression analysis, Procrustes, and Mantel tests were performed to assess the relationships between the biodegradation profile and trophic groups. Mediation linkage was evaluated using the method for multivariate omnibus distance mediation analysis (MODIMA) (Hamidi et al., 2019) to estimate the proportional influence of benthic microbial communities on the direct influence of environmental variables on the biodegradation profile of the 96 organic compounds. In this analysis, the environmental parameters are the independent variable, the biodegradation profile is the dependent variable, and microbial communities act as the mediator. The Euclidean distance of the environmental variables was used as the exposure factor, Bray-Curtis distances of the microbial communities as mediators, and the biodegradation profile based on the $k_{TCC,pH7}$ rate constants as the response (**Supplementary Table S14**).

*Identification of indicator taxa*

A random forest and non-parametric test method based on An et al. (2019) was implemented in the trans_diff() class of the microeco package to identify important taxa based on seasonal occurrence (i.e., seasonal indicator species). The algorithm was performed with seasons as response variables, and the root variables for building a random forest of classification trees were set by default with 1000 grown trees (**Supplementary Table S15**).

#### Supplementary Figures

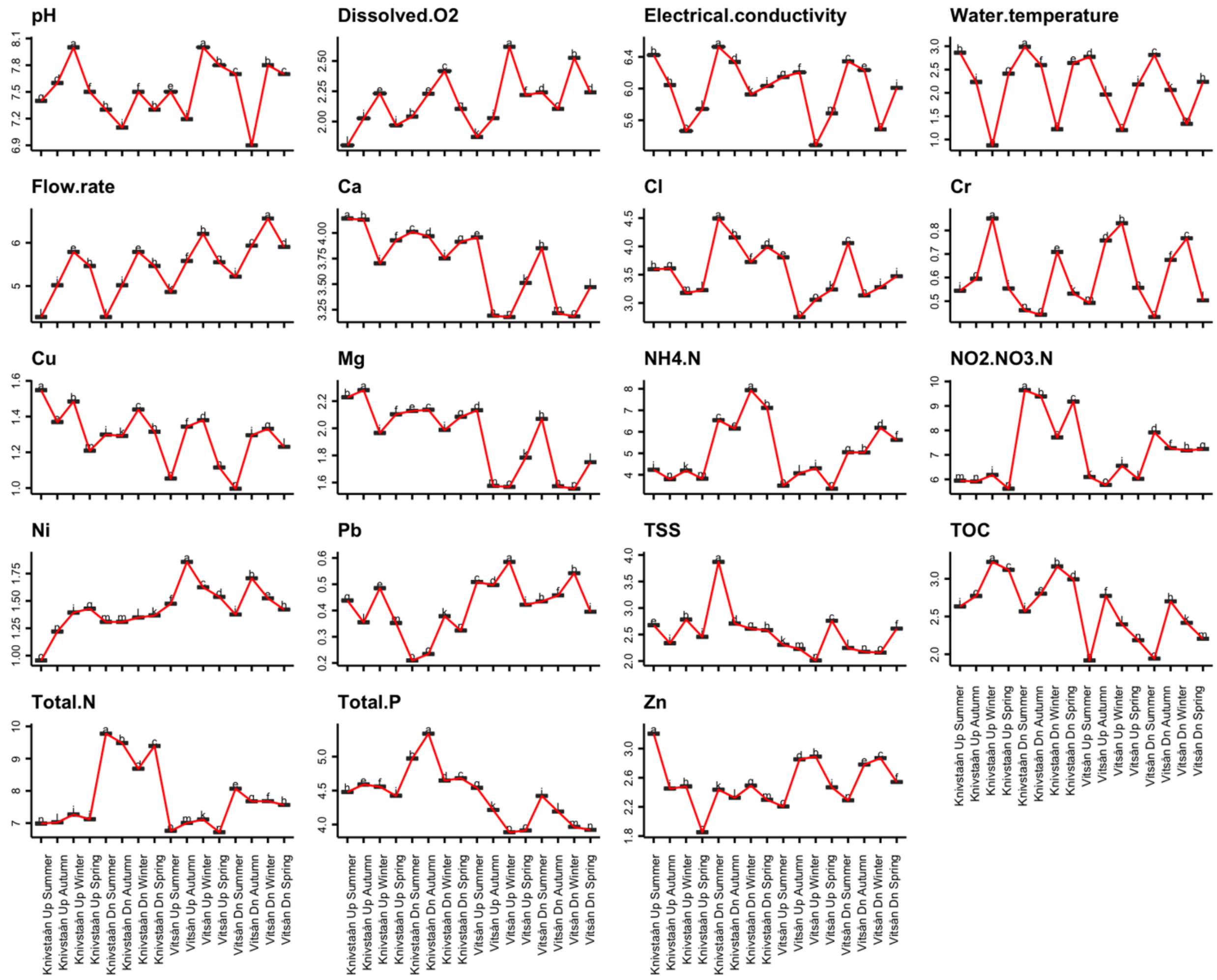


**Supplementary Figure 1.** Visualization of the log10-transformed (except pH) environmental variables grouped by River × Reach × Season: pH, dissolved oxygen (DO, mg/L), electrical conductivity (EC, μS/cm), and water temperature (°C) (Tian et al., 2024). Water chemistry parameters extracted from the national databases: flow rate (L/s), calcium (Ca, mg/L), chloride (Cl, mg/L), chromium (Cr, μg/L), copper (Cu, μg/L), magnesium (Mg, mg/L), ammonium nitrogen ($NH_4$-N, μg/L N), nitrate-nitrite nitrogen ($NO_2$+$NO_3$-N, μg/L), nickel (Ni, μg/L), lead (Pb, μg/L), total suspended solids (TSS, mg/L), total organic carbon (TOC, mg/L), total nitrogen (Total N, μg/L), total phosphorus (Total P, μg/L), and zinc (Zn, μg/L). Different letters indicate differences between groups tested by ANOVA ($p < 0.05$).

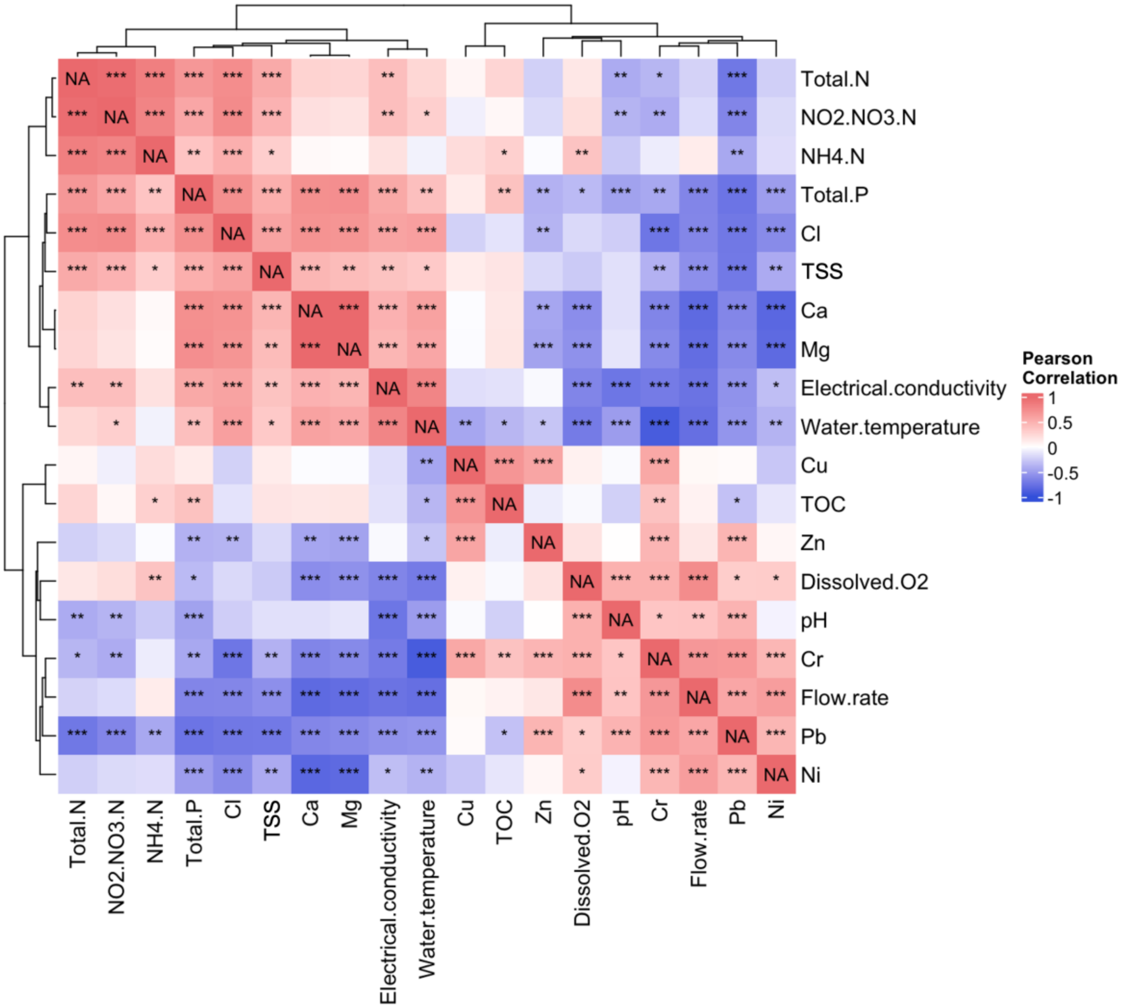


**Supplementary Figure 2.** Pearson's correlation on the relationship between the environmental variables. Positive correlations are indicated in red, and negative correlations in blue. pH, dissolved oxygen (DO, mg/L), electrical conductivity (EC, µS/cm), and water temperature (°C) (Tian et al., 2024). Water chemistry parameters extracted from the national monitoring database: flow rate (L/s), calcium (Ca, mg/L), chloride (Cl, mg/L), chromium (Cr, µg/L), copper (Cu, µg/L), magnesium (Mg, mg/L), ammonium nitrogen ($NH_4$-N, µg/L N), nitrate-nitrite nitrogen ($NO_2$+$NO_3$-N, µg/L), nickel (Ni, µg/L), lead (Pb, µg/L), total suspended solids (TSS, mg/L), total organic carbon (TOC, mg/L), total nitrogen (Total N, µg/L), total phosphorus (Total P, µg/L), and zinc (Zn, µg/L). Different letters indicate differences between groups tested by ANOVA ($p < 0.05$). Hierarchical clustering was performed with the complete linkage method. The asterisk indicates p-values: *** at < 0.001, ** at < 0.01, and * at < 0.05.

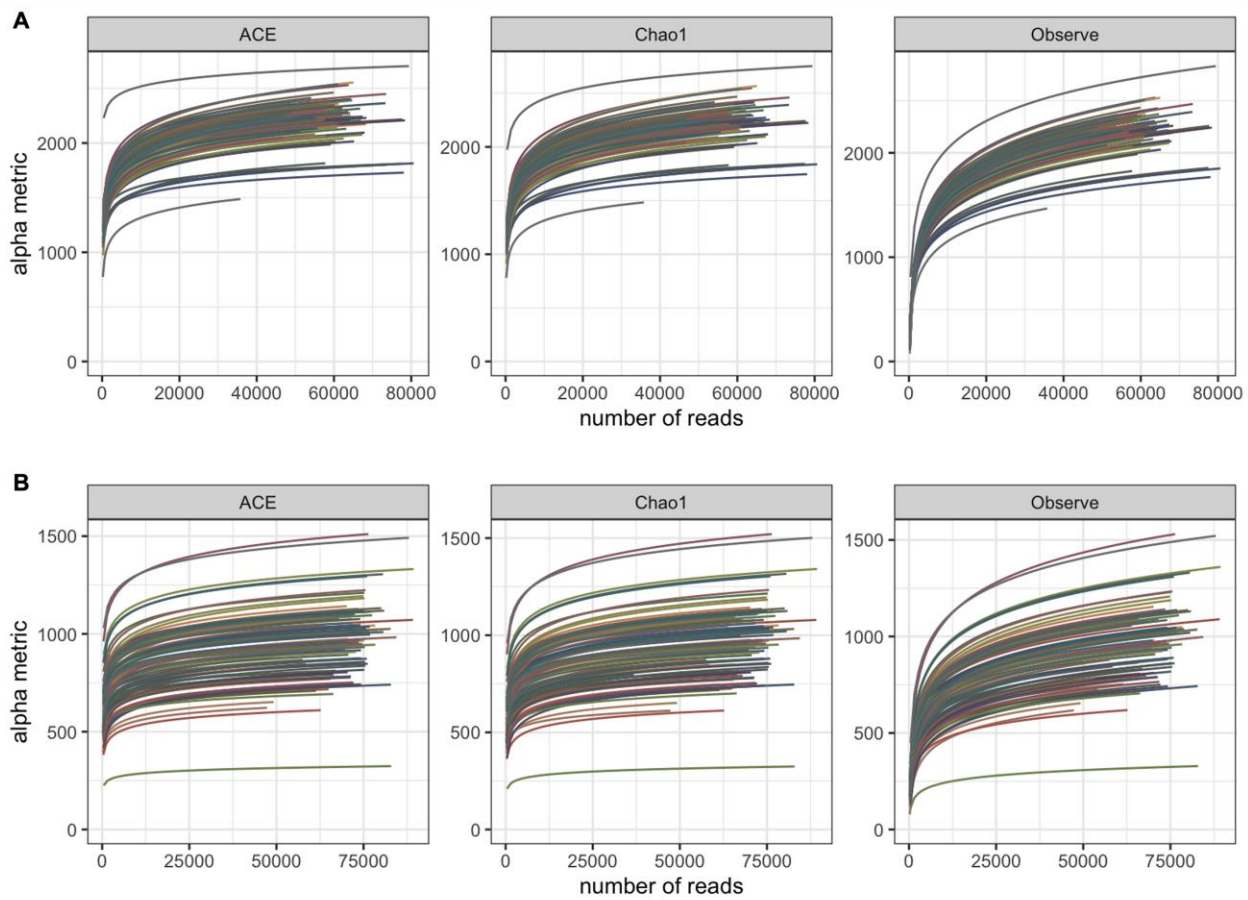


**Supplementary Figure 3.** Rarefaction curves of alpha diversity metrics based on ACE, Chao1, and observed amplicon sequence variants (ASVs) for the (A) 16S rRNA, and (B) 18S rRNA amplicon sequence data. Different colors indicate individual samples.

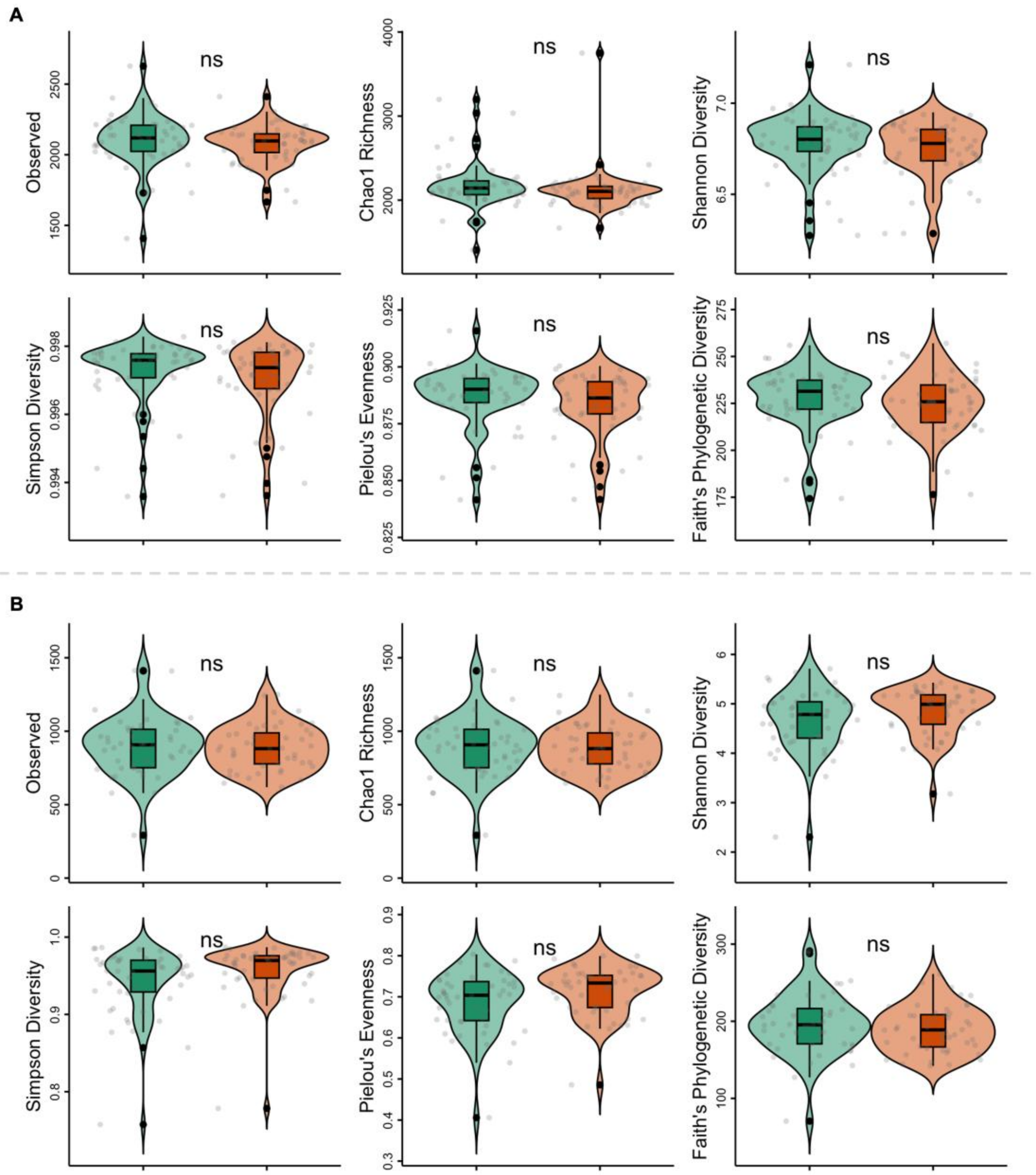


**Supplementary Figure 4.** Alpha diversity estimates of the (A) 16S and (b) 18S data. Green represents the field samples and orange the batch samples. "ns" indicates no significant differences between groups tested by t-test at $p < 0.05$.

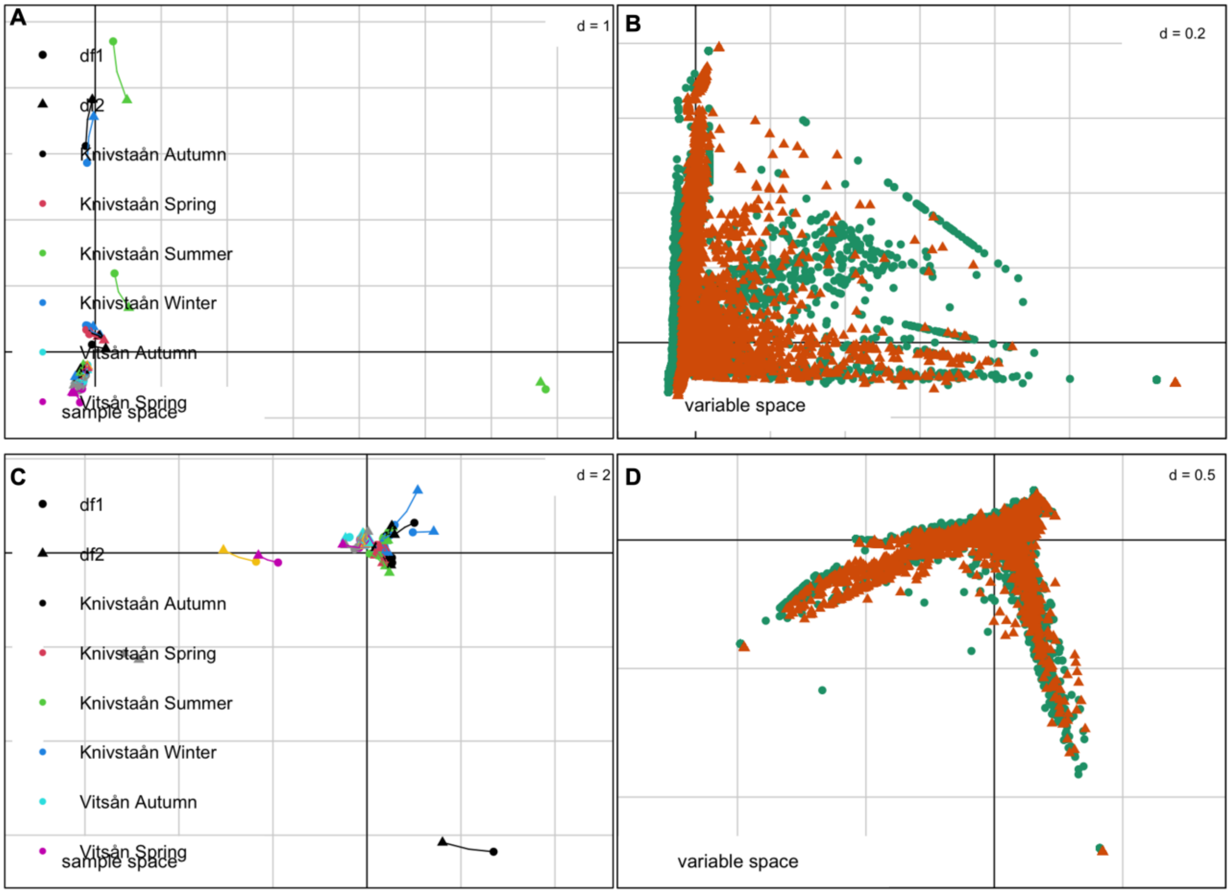


**Supplementary Figure 5.** Co-inertia analysis (CIA) of the field (df1) and batch data (df2). (A) Plot shows the first two components in sample space, and (B) shows the variable space of MCIA for the 16S data, and plots (C-D) for the 18S data, respectively.

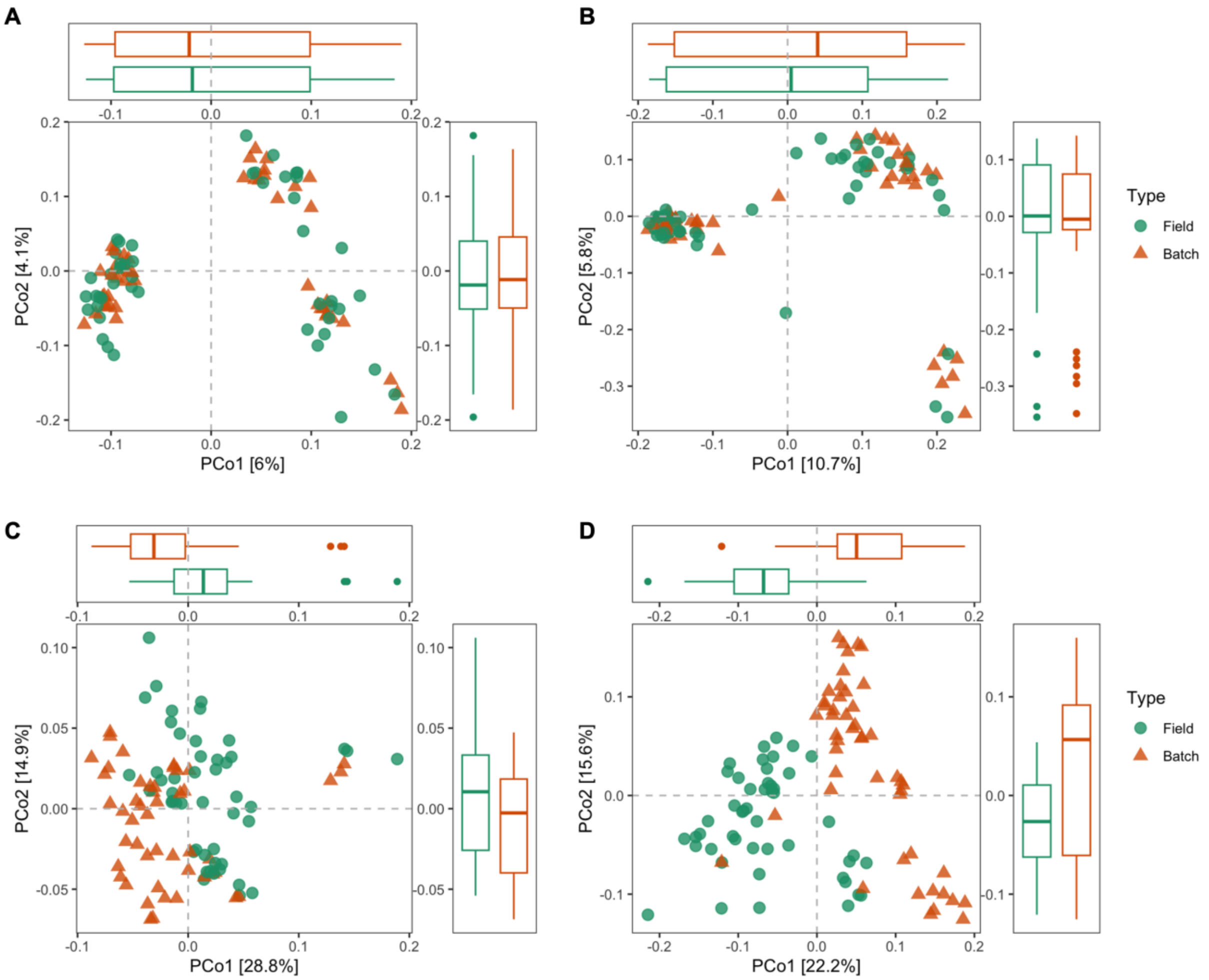


**Supplementary Figure 6.** Principal coordinate analysis (PCoA) plot of the field and batch community dissimilarities based on unweighted UniFrac distance for the (A) 16S and (B) 18S data, and on weighted UniFrac distance for the (C) 16S and (D) 18S data.

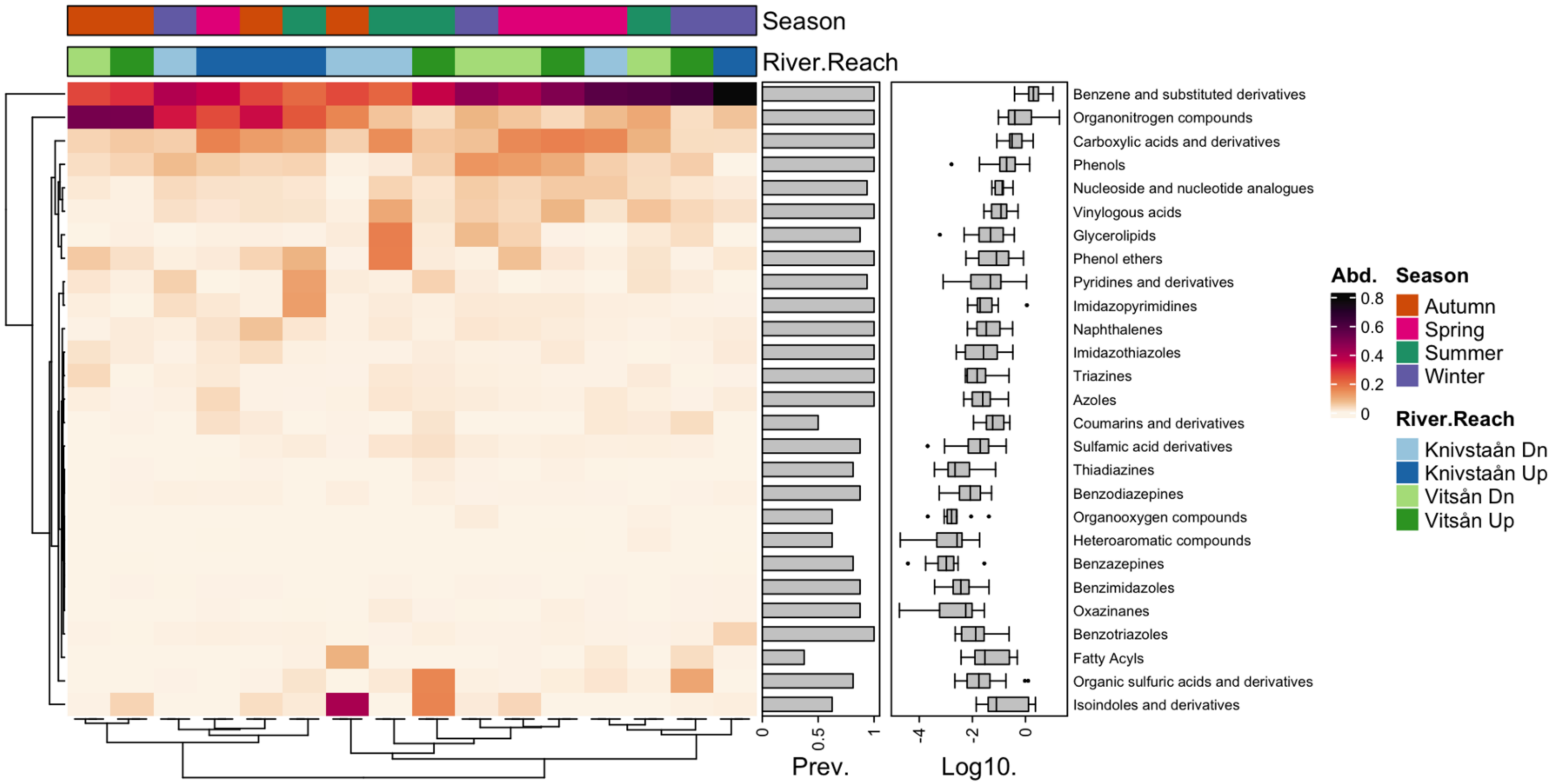


**Supplementary Figure 7.** Spatiotemporal pattern of the biodegradation rates grouped at the class level across the rivers and their reaches. Hierarchical clustering analysis based on Euclidean distance.

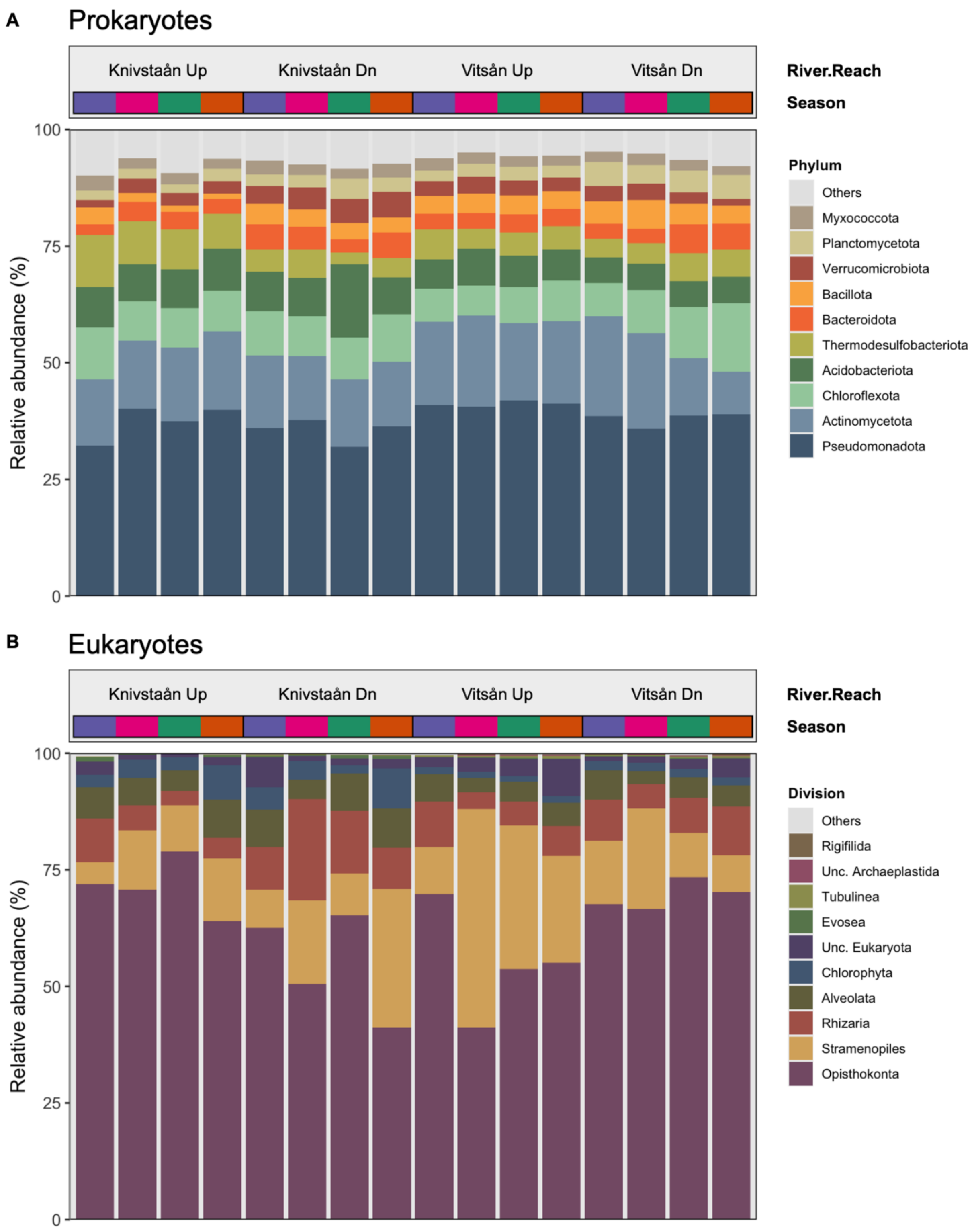


**Supplementary Figure 8.** Relative abundance and composition of the top 10 (A) prokaryotes at the phylum level, and (B) eukaryotes at the division level.

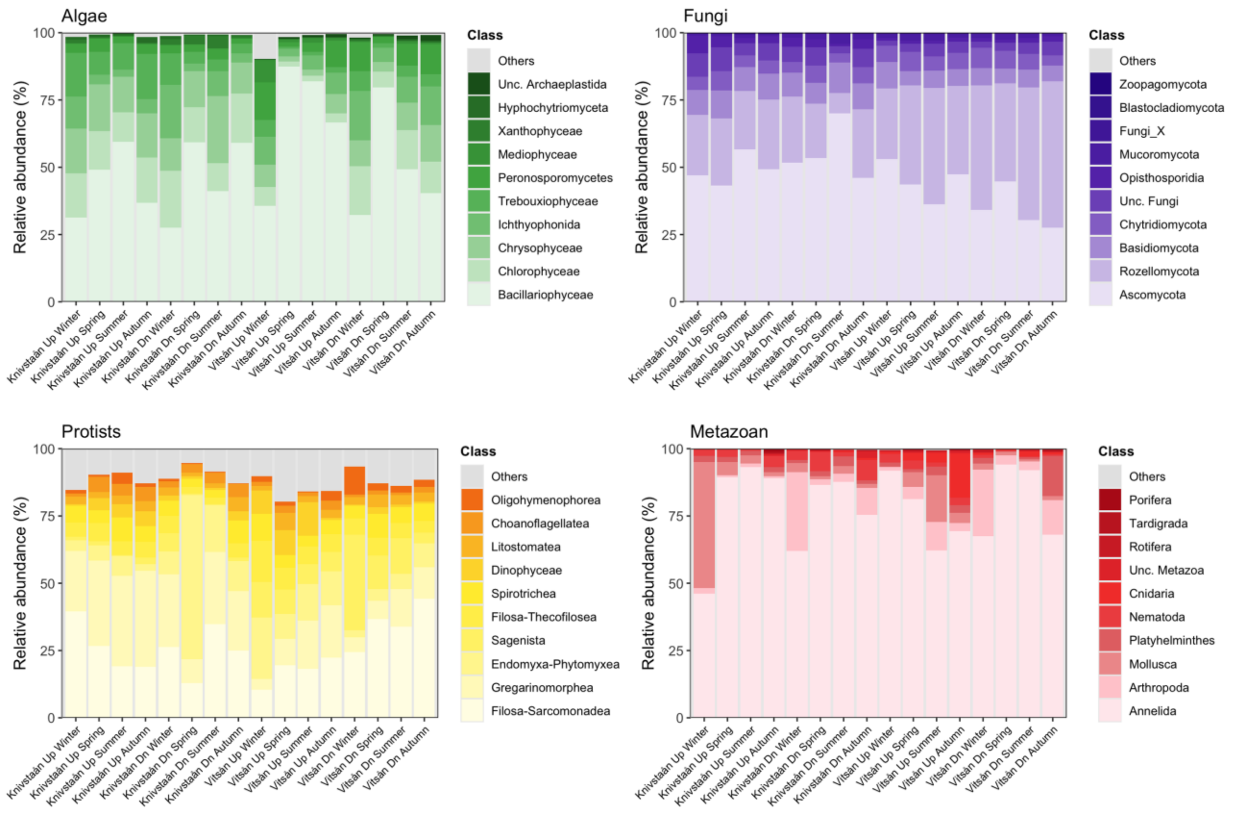


**Supplementary Figure 9.** Relative abundance and composition of the top 10 eukaryotic trophic groups at the class level.

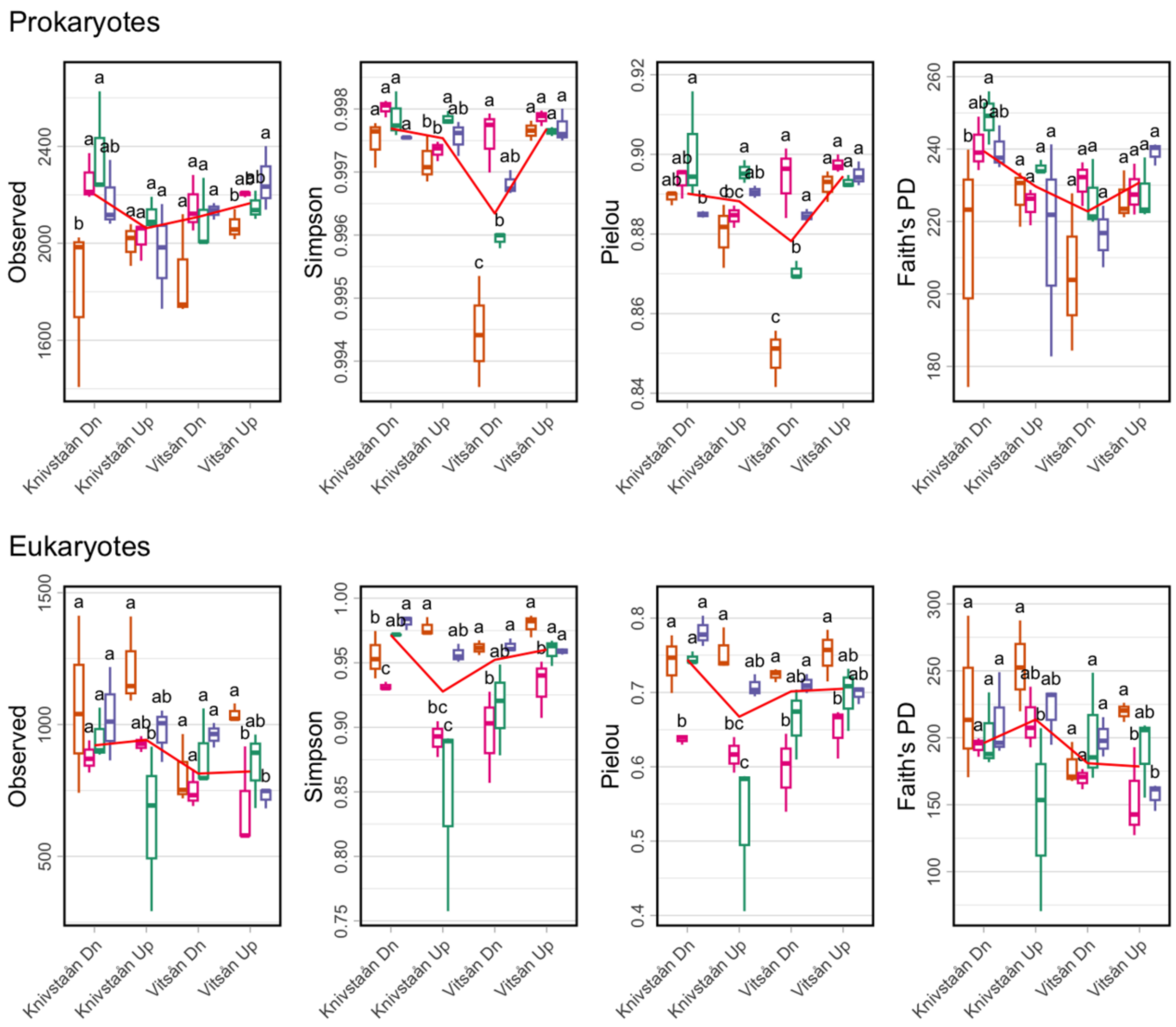


**Supplementary Figure 10 (1 of 2).** Alpha diversity estimates of the prokaryotes and eukaryotes. Different letters indicate differences between groups tested by ANOVA ($p < 0.05$).

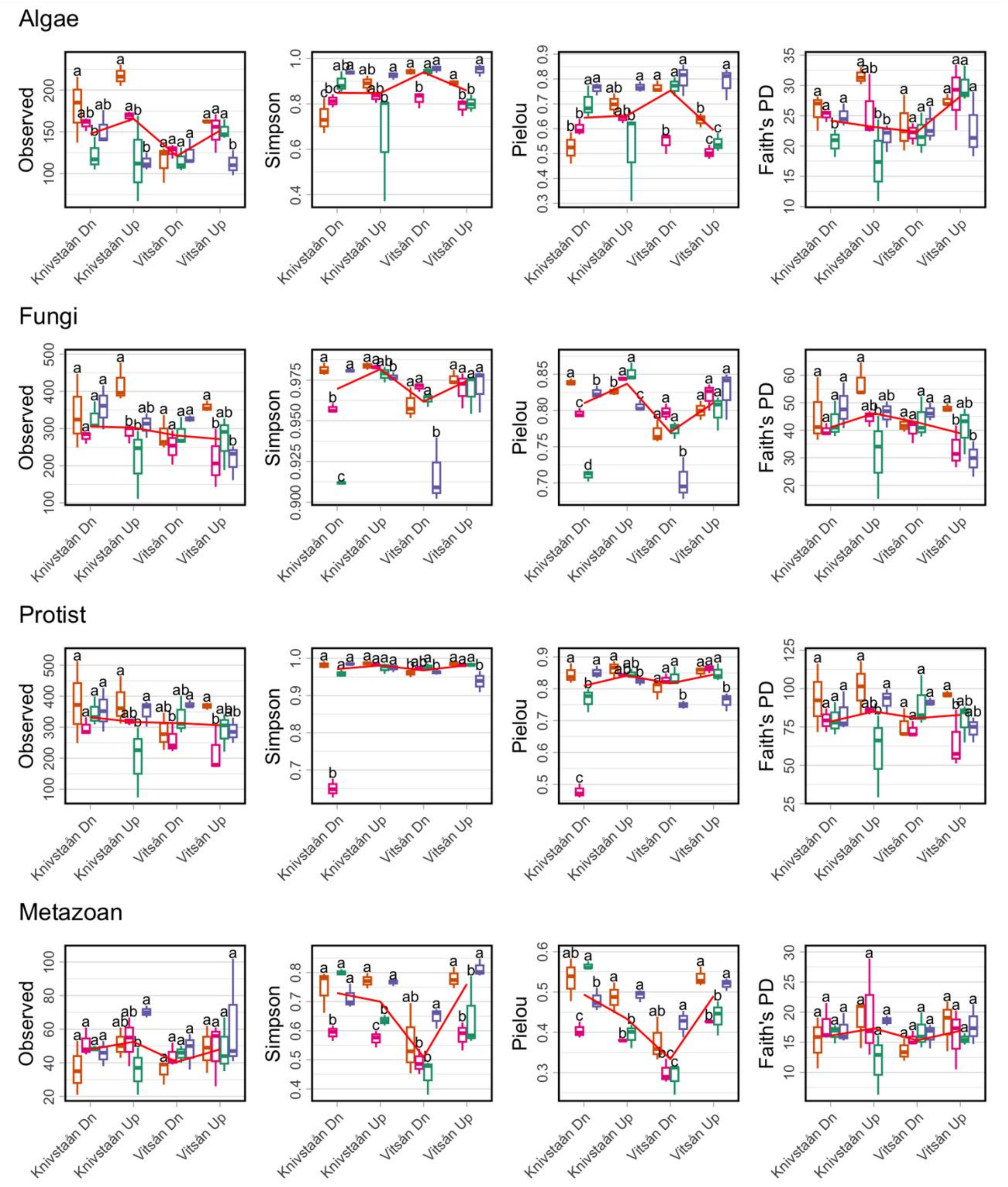


**Supplementary Figure 10 (2 of 2).** Alpha diversity estimates of the eukaryotic groups. Different letters indicate differences between groups tested by ANOVA ($p < 0.05$).

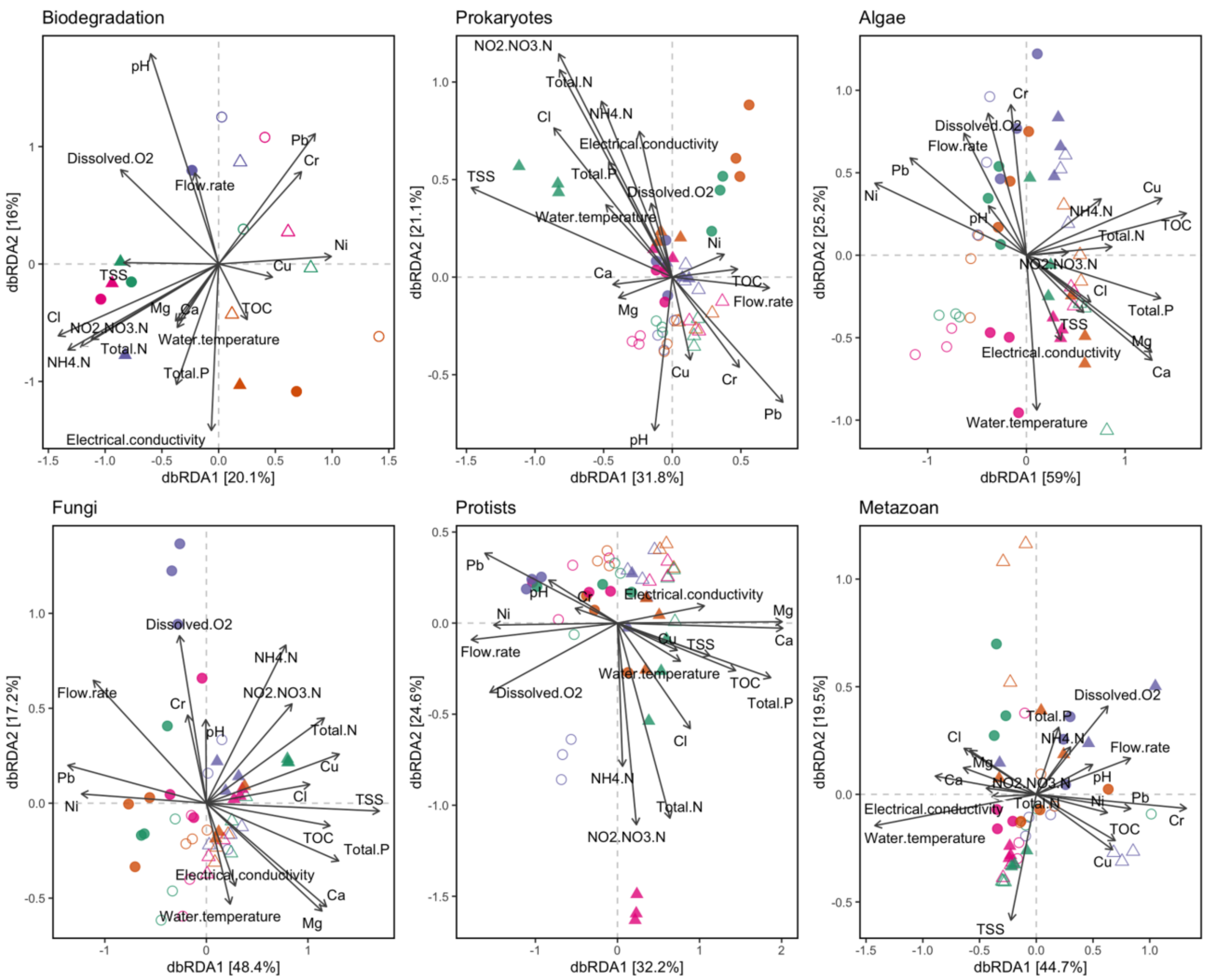


**Supplementary Figure 11.** Distance-based redundancy analysis (dbRDA) of the biodegradation profile based on Bray-Curtis dissimilarity index, and the multi-trophic groups based on weighted UniFrac distance against the environmental factors: pH, dissolved oxygen (DO, mg/L), electrical conductivity (EC, µS/cm), and water temperature (°C) (Tian et al., 2024). Water chemistry parameters extracted from the national monitoring database: flow rate (L/s), calcium (Ca, mg/L), chloride (Cl, mg/L), chromium (Cr, µg/L), copper (Cu, µg/L), magnesium (Mg, mg/L), ammonium nitrogen ($NH_4$-N, µg/L N), nitrate-nitrite nitrogen ($NO_2$+$NO_3$-N, µg/L), nickel (Ni, µg/L), lead (Pb, µg/L), total suspended solids (TSS, mg/L), total organic carbon (TOC, mg/L), total nitrogen (Total N, µg/L), total phosphorus (Total P, µg/L), and zinc (Zn, µg/L).

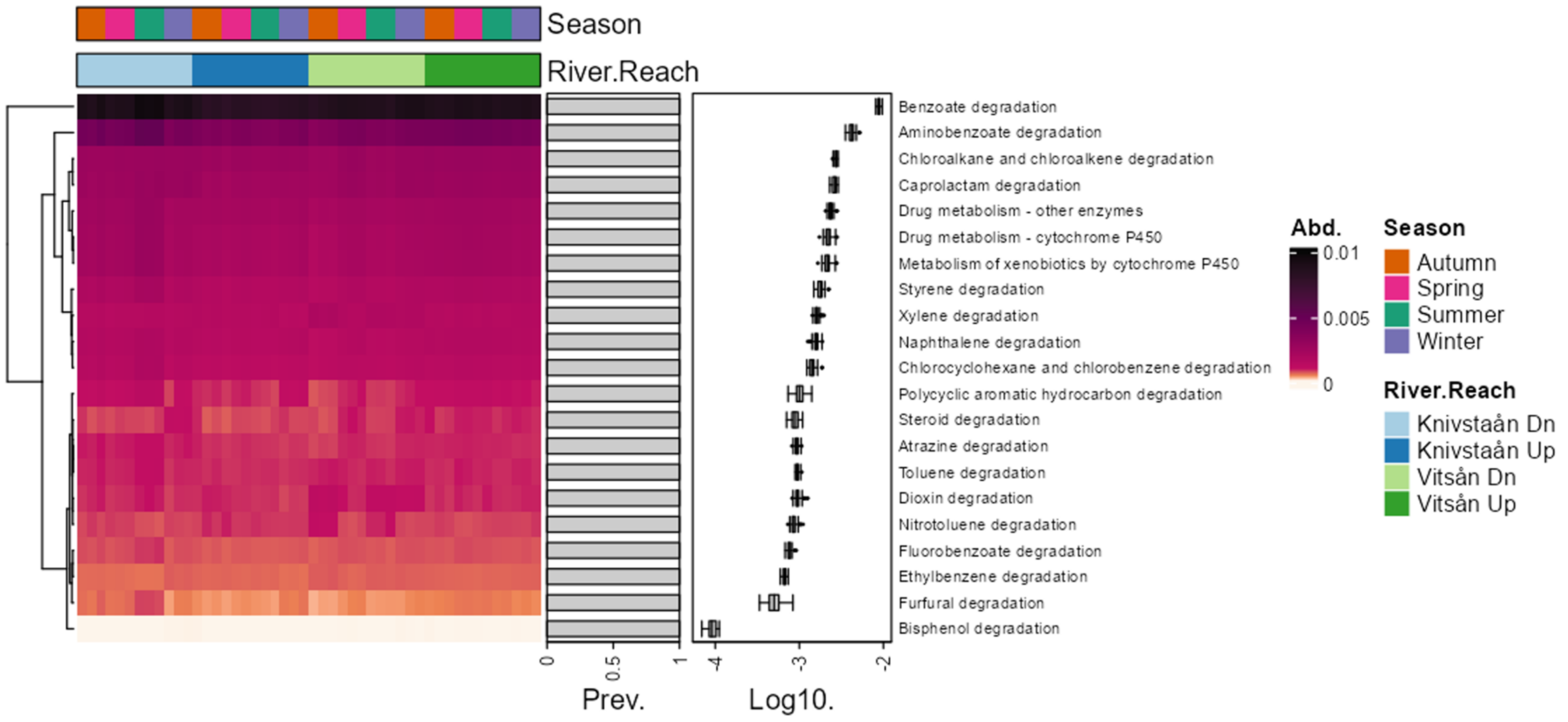


**Supplementary Figure 12.** Heatmap of the predicted functions based on KEGG metabolic pathways (annotations under Level 2 – Xenobiotics biodegradation and metabolism).

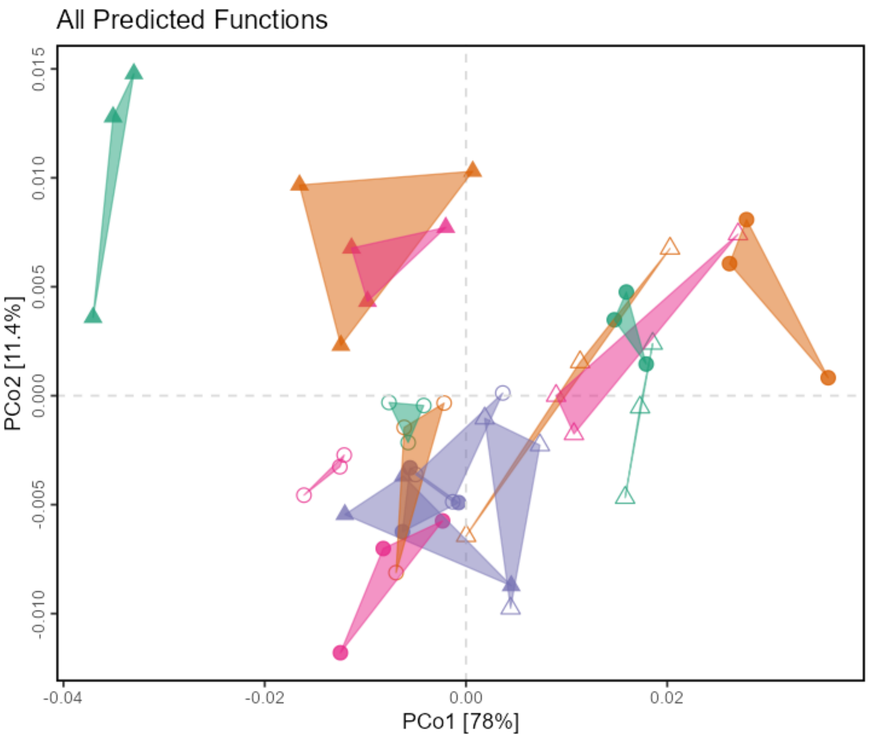


**Supplementary Figure 13.** Principal coordinate analysis (PCoA) based on Bray-Curtis distance of the predicted functions at the spatiotemporal River × Reach × Season grouping.

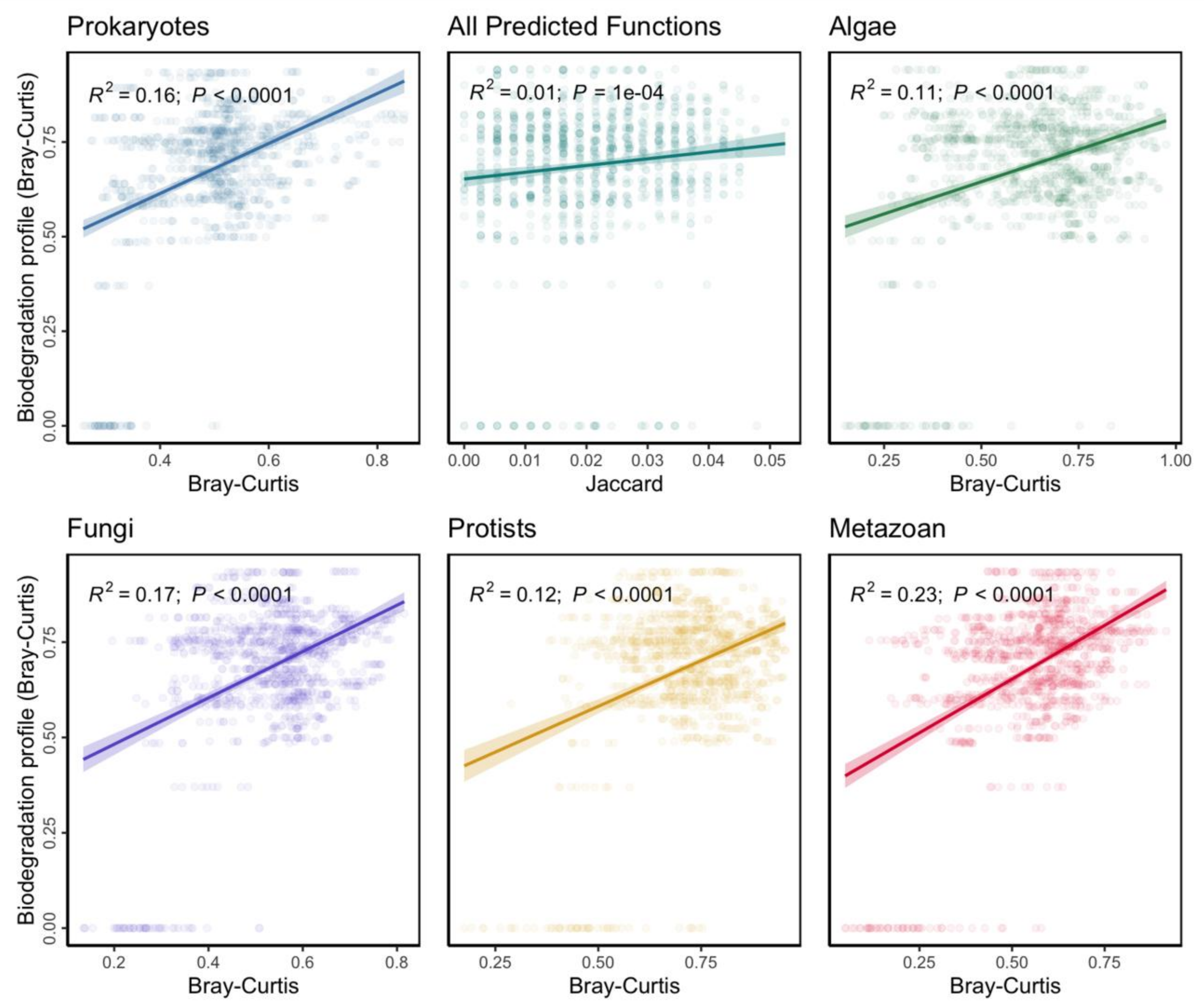


**Supplementary Figure 14.** Correlation between the biodegradation profile and the beta diversity of each multi-trophic group.

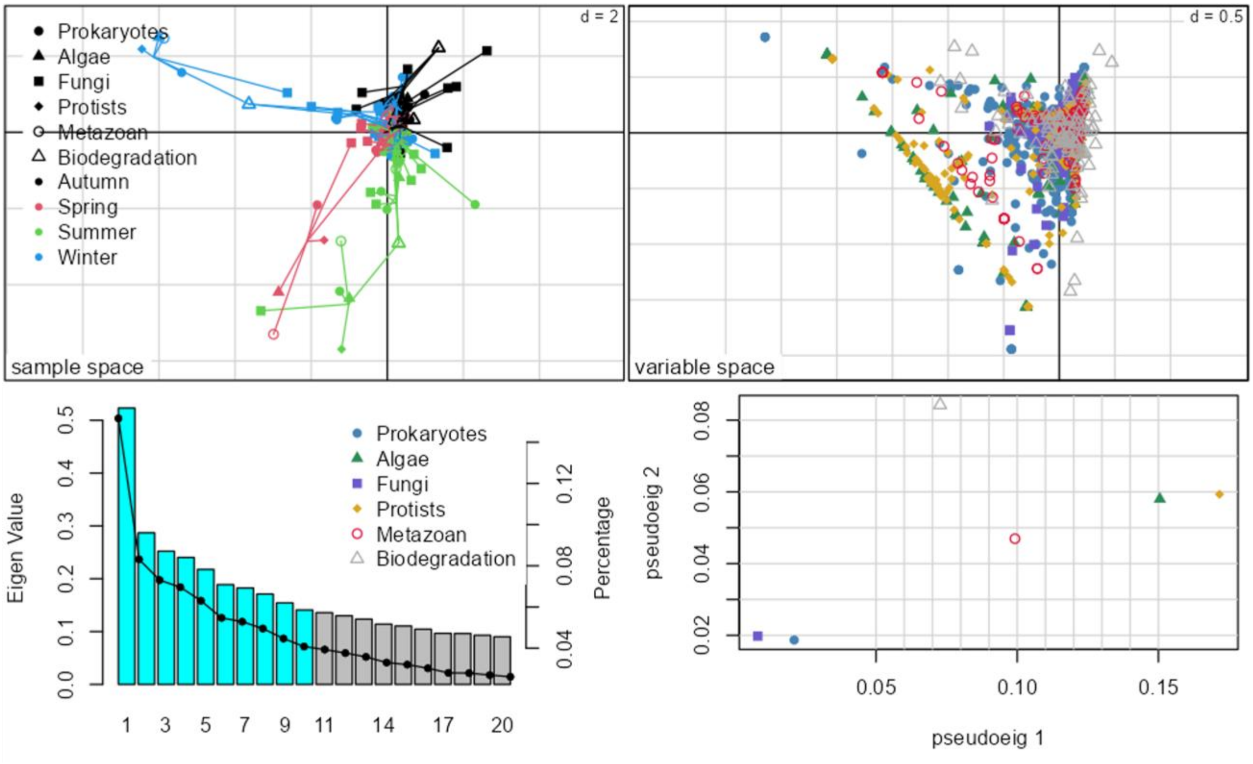


**Supplementary Figure 15.** Multiple CIA (mCIA) analysis of the biodegradation profile and the multitrophic groups. Plot shows the first two components in sample space (top left). Shows the variable space of MCIA (top right). A scree plot of the eigenvalues (bottom left) and a plot of data weighting space (bottom right).

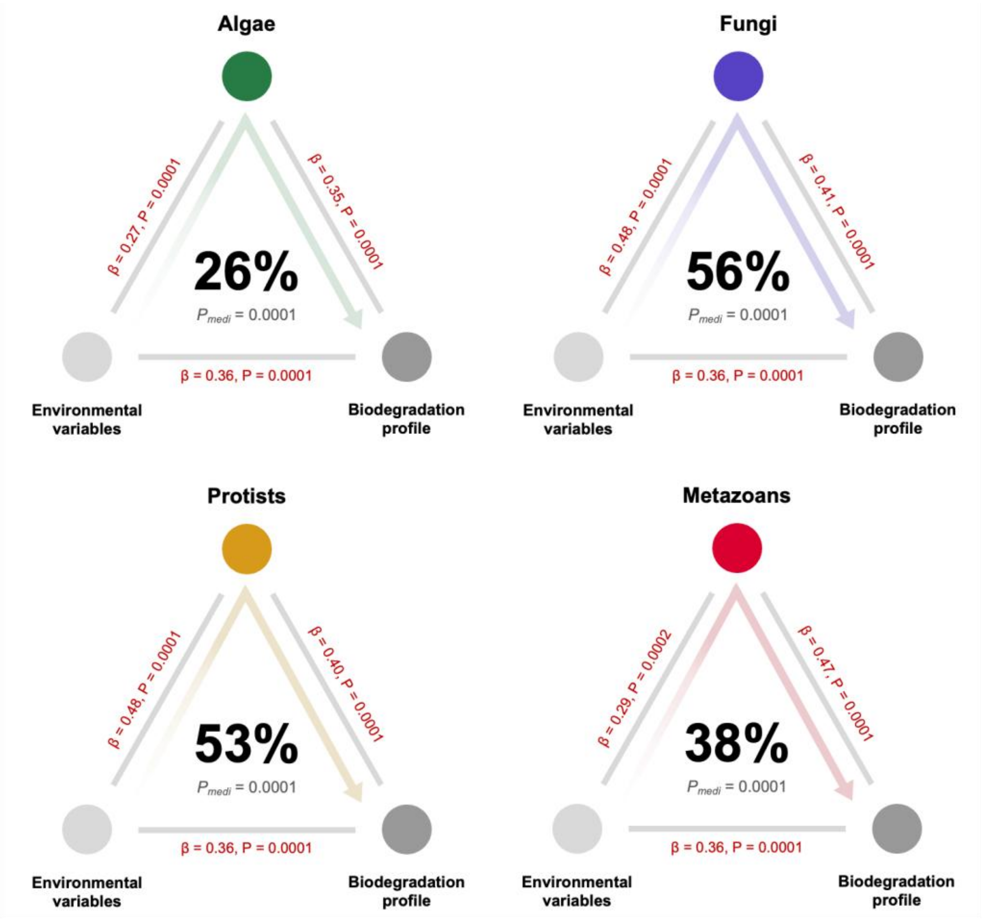


**Supplementary Figure 16.** Mediation association of each eukaryotic trophic group as the mediators of the influence of environmental factors on the biodegradation profile. Beta coefficients and P-values are listed at each connecting line, and the proportions of indirect effect (mediation effect) and mediation P-values (Pmedi) are presented at the center of the ring charts.

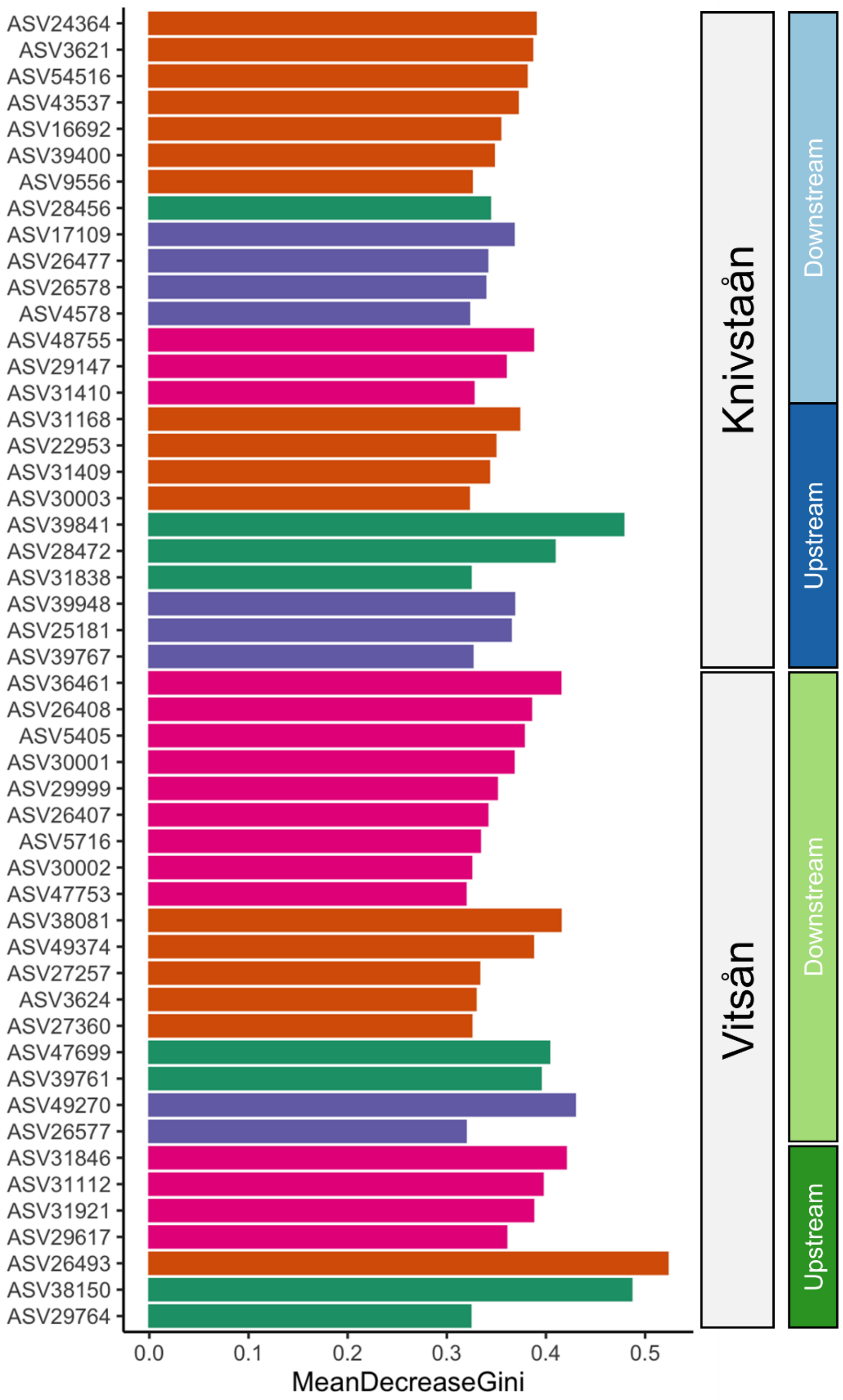


**Supplementary Figure 17.** Differential abundance analysis based on a random forest test on the biodegradation profile of the organic compounds. Only showing top 50 ASVs.

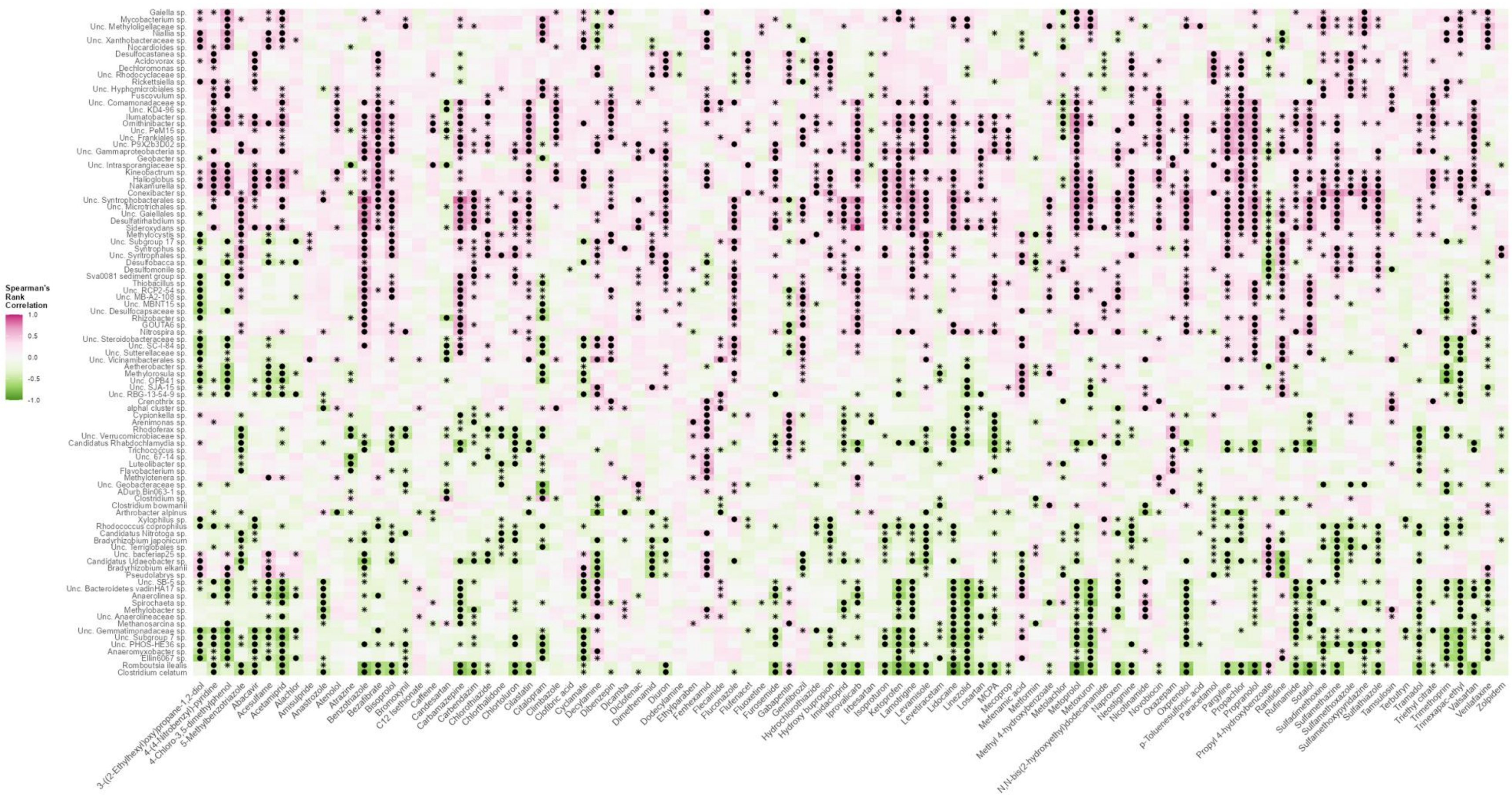


**Supplementary Figure 18.** Heatmap of differentially abundant species across River × Reach × Season group.
